# Machine learning and phylogenetic analysis allow for predicting antibiotic resistance in *M. tuberculosis*

**DOI:** 10.1101/2023.09.06.556328

**Authors:** Alper Yurtseven, Sofia Buyanova, Amay Ajaykumar Agrawal, Olga O. Bochkareva, Olga V. Kalinina

## Abstract

Antimicrobial resistance (AMR) poses a significant global health threat, and an accurate prediction of bacterial resistance patterns is critical for effective treatment and control strategies. In recent years, machine learning (ML) approaches have emerged as powerful tools for analyzing large-scale bacterial AMR data. However, ML methods often ignore evolutionary relationships among bacterial strains, which can greatly impact performance of the ML methods, especially if resistance-associated features are attempted to be detected. Genomewide association studies (GWAS) methods like linear mixed models accounts for the evolutionary relationships in bacteria, but they uncover only highly significant variants which have already been reported in literature. In this work, we introduce a novel phylogeny-related parallelism score (PRPS), which measures whether a certain feature is correlated with the population structure of a set of samples. We demonstrate that PRPS can be used, in combination with SVM- and random forest-based models, to reduce the number of features in the analysis, while simultaneously increasing models’ performance. We applied our pipeline to publicly available AMR data from PATRIC database for *Mycobacterium tuberculosis* against six common antibiotics. Using our pipeline, we re-discovered known resistance-associated mutations as well as new candidate mutations which can be related to resistance and not previously reported in the literature.

## Introduction

*Mycobacterium tuberculosis* (Mtb), the causative agent of tuberculosis (TB), has been a major threat to public health for many years, and remains such a threat now. According to the World Health Organization (WHO), the estimated number of TB-caused deaths in 2021 alone was 1.6 million (1). TB continues to pose a significant threat to global public health because of its ability to easily transmit and the occurrence of drug-resistant strains of Mtb. In 2019, WHO reposterd over ten million cases, including up to 4.5% of infections with drug resistant bacteria (1). Early diagnosis and effective treatment are important steps in controlling the spread of TB and reducing its effect on public health. Given long cultivation time and complex phenotypic resistance testing of Mtb, screening for genetic markers of resistance presents an attractive alternative (2).

Known TB drugs act on Mtb via three mechanisms: first, by preventing the synthesis of the enzymes that makes up the cell wall; second, by interfering with ribosomes that affects protein production; and third, by inhibiting several DNA-level activities, including RNA/DNA synthesis (3). Despite much research conducted on the subject, the drug resistance of Mtb is not fully understood, yet it is known that single nucleotide polymorphisms (SNPs) and other polymorphisms like insertions and deletions (INDELS) play a crucial role in that (4).

The increasing utilization of whole genome sequencing (WGS) for Mtb strains opens up new possibilities in identifying antimicrobial resistance. First, the phylogenetic-based methods such are phylogenetic convergence test and identification of genes under positive selection specific to resistant genomes were successfully applied for hundreds of genomes revealing genes and intergenic regions putatively responsible for resistance (5, 6). Another commonly employed method for detecting significant resistance associated mutations in the data is genome-wide association study (GWAS). For bacteria, information about their population variations is primarily derived from sequences of their genomes. This, in combination with the fact that bacteria have very long genomic segments with strong linkage disequilibrium, creates a very specific setup for bacterial GWAS. This is aggravated by the presence of loci with multiple allelic variants and a large accessory genome (genes that are present only in some strains of a bacterial species, but not in others) (7). Indeed, in Mtb recombination rates are particularly low (8), and thus virtually all loci of the genome are in linkage disequilibrium. On the other hand, the accessory genome of Mtb is very small, in contrast to other bacterial species, in which genes conferring resistance to particular antibiotics are often transferred via plasmids (9, 10).

Strong linkage disequilibrium between loci in bacteria implies that the population structure plays a major role and should be accounted for in GWAS studies. Indeed, many passenger mutations may be associated with a phenotype-relevant variant and will be called together with it, because they all step from a branch of closely related strains on the species’ phylogenetic tree. Such effects have been accounted for by using linear mixed models . In these models, the effect of each locus on the phenotype is modeled in the context of all other loci that are considered to contribute random effects. In this way, the effect of each locus that is strongly correlated with the background is systematically decreased. Linear mixed models showed promising results in bacterial GWAS for resistance phenotypes in many species including *E. coli, S. aureus, K. pneumoniae*, and Mtb (11). Combination of GWAS approach with a phylogenetic convergence test in Mtb significantly improved the approach and allowed to identify epistatic interactions between drug-resistance-associated genes (12). This approach is based on the idea that the true resistance-conferring mutations often originate at multiple branches of the phylogenetic tree of Mtb strains, while non-relevant passenger mutations occur in single (but maybe very populated) branches. Such effects are not visible when the strains are considered to be independent as in classical GWAS, but may be accounted for when population structure is taken into account.

Apart from population structure and epistasis, other factors like recombination rate, within-host diversity, polygenicity and multi-allelic SNPs also need to be taken into account while performing the GWAS in bacteria. Various computational tools and methods have been developed to account for these factors (13). For example, CCTSWEEP (14), VENN (14) and GWAMAR (15), use phylogenetic trees to account for population structure, but do not take other factors into account. Other phylogenetic tree based methods like treeWAS (16) takes all factors into account except within-host diversity, while Scoary (17) does not consider within-host diversity as well as recombination rate. Among all the available bacterial GWAS tools, SEER (18) and pyseer (python implementation of SEER) (19) are the only two that considers most of the pitfalls one can stumble upon in bacterial GWAS.

Over the last few years, certain rule-based approaches like TB-Profiler (20) have been developed to detect the phenotype of newly sequenced Mtb strains. These approaches work by calling out the variants and comparing them against curated databases. Thus, these approaches can only detect resistance caused by known markers. To overcome this, prediction approaches based on machine learning (ML) have been recently explored for the identification of resistance associated mutations in bacteria. For example, 414 strains of *P. aeruginosa* have been investigated with a host of classical machine-learning techniques while accounting for population structure with a training strategy based on blocks of phylogenetically related sequences (21). Machine learning has been used for detecting resistance in Mtb data as well, employing different ML methods and achieved area under ROC curve values up to 0.95 in a classification task for resistance towards selected drugs using features from 23 selected target genes known to be implicated in resistance development (22). A recent computational framework, TB-ML, provides implementations for different ML methods such are random forest, direct association and convolutional neural networks (23). Training datasets for these methods can be found in public resources, such as, for example, the PATRIC database (24).

However, to date most models trained with ML algorithms do not account for specifics of the bacteria data, such as population structure and linkage disequilibrium, and thus can be prone to misinterpretation. Therefore, just as for GWAS studies, ranking genetic variants is a crucial part that should be conducted before applying algorithms for model training to filter out non-significant features from training dataset. One of suggested ways to rank such variants is to predict their potential impact on protein function (25). An indirect way to take population structure into account is to design the training process specifically for each set of bacterial genomes by splitting the isolates into training, test, and validation sets based on genomic distance between them (26).

In this study, we present a novel phylogeny-based method for ranking genetic variants followed by training ML models for predicting antibiotic resistance in Mtb. We demonstrate that this filtering is crucial and improves performance of the ML models. Using bacterial GWAS methods as a baseline, we identified known resistance-associated variants in a set of Mtb strains with known resistance profiles from the PATRIC database, as well as detect novel potential resistance-associated variants.

## Materials and Methods

### Genomic and phenotypic data and variant calling

Genome sequences and the resistant phenotype data of the Mtb strains for six antibiotics (amikacin, capreomycin, ethionamide, kanamycin, ofloxacin and streptomycin) were downloaded from PATRIC database (24) (retrieved on November 26, 2021). First, the strains were filtered out because the corresponding fasta files were corrupted in the database. Next, the duplicate genomes and the cases when identical genomes were annotated with conflicting phenotypes were further removed. Additionally, genomes with more than 5 consecutive ‘N’ nucleotides, L90 > 100 and those comprising more than 999 contigs were excluded from our study. To get rid of contamination, all genomes with maximum pairwise Mash genetic distance exceeding 0.2 to any other genome were removed. Finally, genomes with the length deviating by more than two standard deviations from the average Mtb genome length and those with more than 5% contamination were excluded from our study. After applying these filters, our final dataset consisted of 4869 genomes (Suppl. Table 6) which were used in our study (Table 1).

**Table 1.** Datasets used in this study. For the number of strains and SNPs, the final numbers after all filtering are provided.

| Drug name | Line of therapy | Pharmacological group | Number of strains | Number (fraction) of resistant strains | Number of SNPs (features) |
| --- | --- | --- | --- | --- | --- |
| Streptomycin | First line | Aminoglycosides | 4,726 | 1158 (24,5%) | 24,425 |
| Amikacin | Second line | Aminoglycosides | 1,149 | 208 (18,1%) | 18,864 |
| Capreomycin | Second line | Aminoglycosides | 1,086 | 205 (18,9%) | 17,045 |
| Kanamycin | Second line | Aminoglycosides | 1,362 | 297 (21,8%) | 17,335 |
| Ofloxacin | Second line | Fluoroquinolones | 795 | 307 (38,6%) | 14,185 |
| Ethionamide | Second line | Nicotinamide derivative | 571 | 210 (36,8%) | 12,974 |
| Total |  |  | 4,869 |  | 24,425 |

Whole genome sequences of all the 4869 strains were mapped to the H37Rv reference genome (NCBI accession: NC_000962) and variant calling was performed using the Snippy (27). Variants present in < 0.2% strains were filtered out to account for sequencing errors. Among the variants, we removed INDELS and only retained single-nucleotide polymorphism (SNPs). Finally, we were left with 24,425 SNPs which were then used as a features for further downstream studies.

### Genome wide association studies (GWAS)

As a baseline, genome wide association analysis was carried out using pyseer (19), since this tool accounts for most of the pitfalls that one can come across in bacterial GWAS (13). To account for strong population structure, a similarity kinship matrix was generated using “similarity_pyseer” command of pyseer and was used as input for GWAS analysis. Further, the linear mixed models (LMM) which models other SNPs as random effects to control for population structure was used to perform the GWAS analysis. Finally, to control for multiple testing, the Bonferroni correction was used, and a p-value threshold of 0.05 divided by number of variants was adopted for all antibiotics to select the significant variants associated with resistance phenotype.

### Phylogenetic reconstruction and phylogeny-related parallelism score

To build the phylogeny of the Mtb strains, we used the PanACoTA pipeline (28). Orthologous groups were constructed with 80% threshold for protein identity, concatenated codon alignment of 161 single-copies common genes was used to construct the maximum-likelihood phylogenetic tree using fasttree (29) with 1000 bootstrap replicates. The phylogenetic tree with resistance profiles was visualized using the iTOL online tool (30). To calculate the phylogeny-related parallelism score (PRPS) for each SNP we perform the following procedure (adopted from (31)). First we generate a matrix of the pairwise distances between all the nodes in the tree. Then we collapse the clades where all leaves have a SNP and assign this SNP to the corresponding ancestral node. Finally we calculate the PRPS as a logarithm of a sum of pairwise distances between the nodes with SNP:

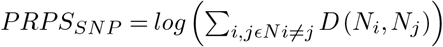

where N are nodes which the SNP is assigned to, D is the phylogenetic distance between them. Thus, PRPS reflects the number of occurrences of a SNP across the tree and distances between the nodes where it occurred.

### Machine learning for predicting antimicrobial resistance

Resistance profiles from the PATRIC database were used as the target variable for machine learning methods. To this end, the minimal inhibitory concentration (MIC) values were converted into a binary variable representing ‘resistant’ and ‘susceptible’ phenotypes. We used two supervised learning models; support vector machine (SVM) with linear kernels and random forest (RF) algorithms. Hyperparameters were optimized with AutoML. All ML models were implemented with scikit-learn (32) and auto-sklearn (33) in Python.

Three feature sets were used for our ML models: (1) all SNPs selected as described above; (2) 30% features with highest PRPS; (3) 30% features chosen randomly from all features. The 30% threshold was chosen, since it consistently delivered best performance for the PRPS-ranked features (data not shown). Further, feature importance analysis was performed to extract predictive resistance genetic markers. For RF models, we used permutation-based feature importance analysis (32). For SVMs we used both permutation-based feature importance analysis and coefficients.

## Results

First, we established a baseline with GWAS analysis. The pyseer software identified key known resistance mechanisms for the corresponding antibiotics (Figure 1, Supp. Table S2): mutations in 16S ribosomal RNA gene for aminoglycosides, mutations in 30S ribosomal protein S12, catalase peroxidase *katG*, and 3-oxoacyl-ACP reductase *fabG* genes for streptomycin, SNPs upstream of the aminoglycoside acetyltransferase *eis* gene for kanamycin, and mutations in DNA gyrase subunit A for ofloxacin. We did not observe any mutations known to confer resistance to ethionamide; instead for this drug, which is a second-line therapyone, we observed a mutation in the 16S ribosomal RNA gene, which may be an indication of multiresistance against first-line streptomycin or other aminoglycosides. In addition, we observed several unreported mutations in and upstream of genes which encode hypothetical proteins and a transcription regulator from the AraX/XylS family that have less significant p-values.

**Fig. 1.**
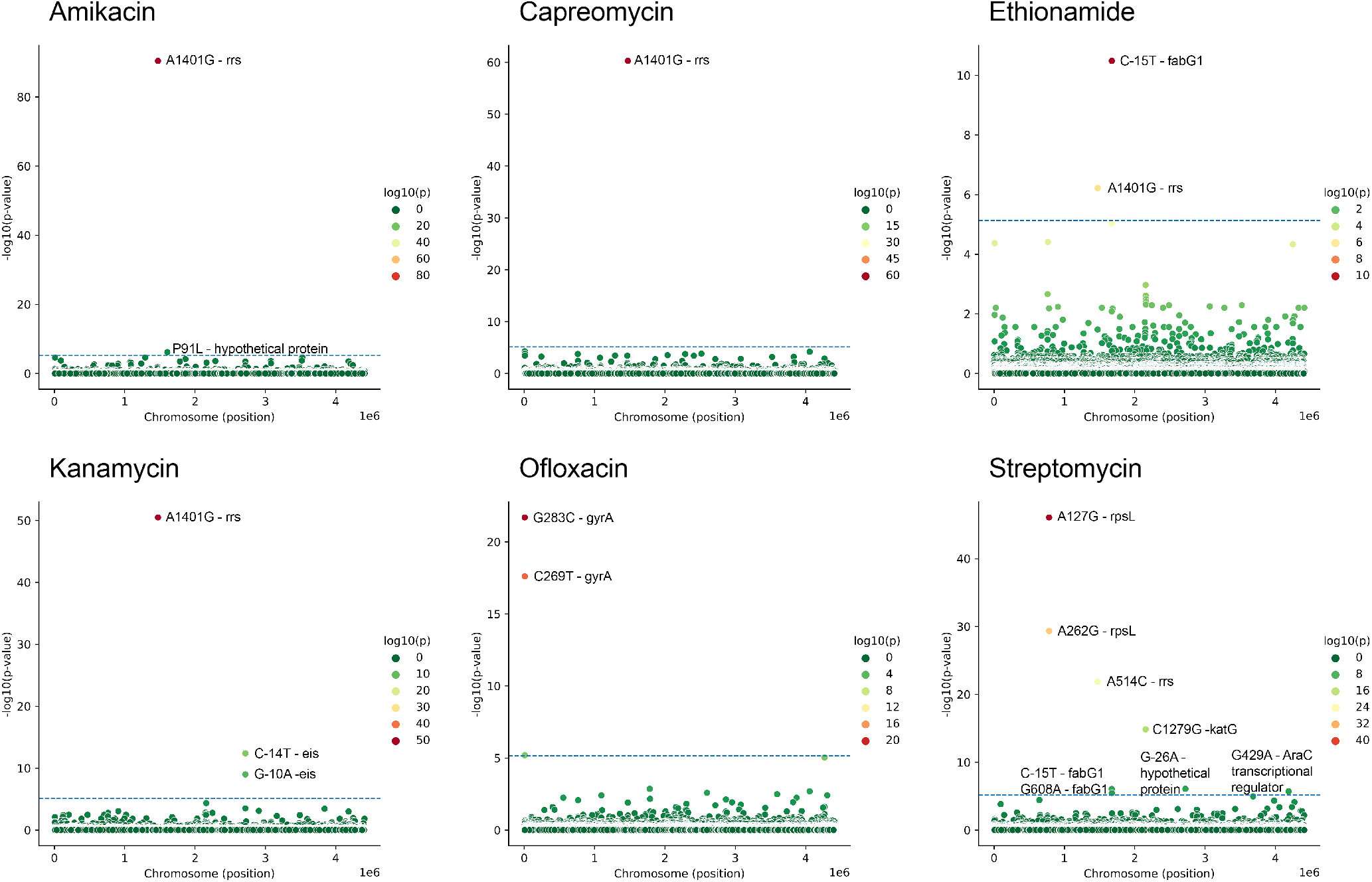
Associations between SNP and antibiotic resistance found using GWAS (see Methods). Horizontal line shows selected threshold for significance.

In the ML analysis, first we calculate phylogeny-related parallelism score (PRPS), a measure of inconsistency between SNP phyletic pattern and the species tree topology, to exclude mutations that are strongly linked with the population structure from the training dataset (see Methods). According to our procedure, SNPs whose distribution is consistent with phylogenetic tree structure have low PRPS. In contrast, high PRPS indicate independent acquisition of SNPs by different lineages. PRPS reflects whether a SNP is monophyletic or polyphyletic, where high PRPS scores correspond to highly polyphyletic features (Figure 2, top). For comparison, a known resistance-associated mutation A90V in GyrA that confers a strong resistance to fluoroquinolones (34), has a PRPS score of 2.824321 and is in the top 11% of the PRPS-ranked feature list (Figure 2, bottom).

**Fig. 2.**
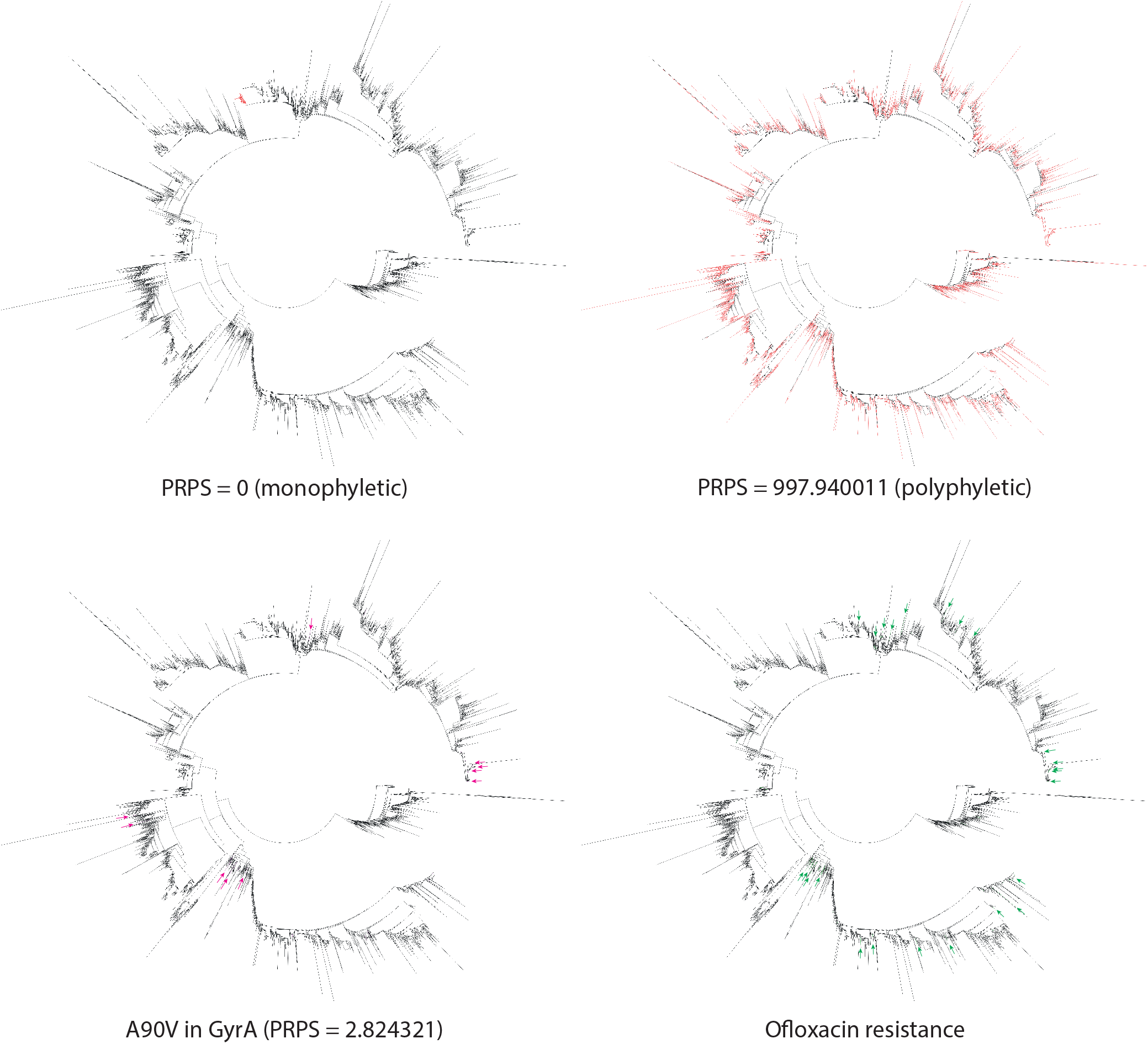
Top: High- and low-PRPS SNPs. Branches corresponding to the strains containing the SNP are colored red. Bottom: Mutation A90V in GyrA associated with resistance to fluoroquinolones (left) and ofloxacin-resistant strains (right). Branches corresponding to strains with the mutation A90V in GyrA are colored magenta and indicated with arrows. Branches corresponding to ofloxacin-resistant strains are colored green, and the corresponding clades are marked with green arrows.

Top 30% of features with highest PRPS were used for training our ML models. In all cases, even though a significant number of mutations (70%) were not considered in the models, both MCC and ROCAUC values increase in almost all cases when high-PRPS mutations are used as features (the only exception being the RF models for amikacin, where the performance is almost perfect anyway). To test the significance of this observation, we randomly deleted 70% of the mutations from our data and re-trained the models. In this case, all performance measures dropped drastically, which proved that SNPs selected by the phylogeny analysis were not random and were enriched in predictive features, while containing less noise (Table 2). Moreover, using high-PRPS features increased the training speed and reduced the amount of resources used for model training in terms of memory and CPU time. Additionally, we checked for presence of known antibiotics resistance markers among the predictive features using features importance analysis, and in all cases these markers are retained in the high-PRPS models and disappear in random-30% models (data not shown). Thus, we used top 30% high-PRPS features for all further analysis.

**Table 2.** Performance of ML models. MCC (Matthew’s correlation coefficient) values of trained models

|  | Support Vector Machine (SVM) |  |  | Random Forest |  |  |
| --- | --- | --- | --- | --- | --- | --- |
| MCC | All features | 30% PRPS | 30% Random | All features | 30% PRPS | 30% Random |
| Amikacin | 0.720 | 0.752 | 0.302 | 0.881 | 0.883 | 0.481 |
| Capreomycin | 0.539 | 0.620 | 0.334 | 0.779 | 0.780 | 0.569 |
| Ethionamide | 0.325 | 0.370 | 0.269 | 0.550 | 0.605 | 0.488 |
| Kanamycin | 0.766 | 0.685 | 0.415 | 0.812 | 0.856 | 0.546 |
| Ofloxacin | 0.508 | 0.549 | 0.294 | 0.778 | 0.778 | 0.452 |
| Streptomycin | 0.602 | 0.613 | 0.477 | 0.782 | 0.801 | 0.650 |
| ROCAUC | All features | 30% PRPS | 30% Random | All features | 30% PRPS | 30% Random |
| Amikacin | 0.842 | 0.872 | 0.663 | 0.927 | 0.918 | 0.765 |
| Capreomycin | 0.779 | 0.816 | 0.683 | 0.858 | 0.863 | 0.718 |
| Ethionamide | 0.665 | 0.686 | 0.640 | 0.780 | 0.807 | 0.746 |
| Kanamycin | 0.873 | 0.844 | 0.718 | 0.880 | 0.916 | 0.741 |
| Ofloxacin | 0.757 | 0.781 | 0.649 | 0.870 | 0.870 | 0.723 |
| Streptomycin | 0.798 | 0.816 | 0.747 | 0.865 | 0.890 | 0.785 |
| Sensitivity | All features | 30% PRPS | 30% Random | All features | 30% PRPS | 30% Random |
| Amikacin | 0.721 | 0.786 | 0.485 | 0.868 | 0.843 | 0.671 |
| Capreomycin | 0.650 | 0.720 | 0.517 | 0.733 | 0.760 | 0.453 |
| Ethionamide | 0.603 | 0.603 | 0.676 | 0.689 | 0.779 | 0.691 |
| Kanamycin | 0.787 | 0.755 | 0.585 | 0.777 | 0.681 | 0.532 |
| Ofloxacin | 0.719 | 0.771 | 0.573 | 0.771 | 0.771 | 0.625 |
| Streptomycin | 0.687 | 0.741 | 0.642 | 0.752 | 0.815 | 0.601 |
| Specificity | All features | 30% PRPS | 30% Random | All features | 30% PRPS | 30% Random |
| Amikacin | 0.965 | 0.958 | 0.840 | 0.987 | 0.994 | 0.858 |
| Capreomycin | 0.910 | 0.912 | 0.849 | 0.983 | 0.979 | 0.982 |
| Ethionamide | 0.727 | 0.769 | 0.603 | 0.810 | 0.835 | 0.802 |
| Kanamycin | 0.961 | 0.933 | 0.851 | 0.983 | 0.992 | 0.949 |
| Ofloxacin | 0.796 | 0.790 | 0.725 | 0.970 | 0.970 | 0.820 |
| Streptomycin | 0.910 | 0.891 | 0.853 | 0.977 | 0.965 | 0.969 |

We further investigated our SVM and RF models with permutation-based feature importance analysis (35) to identify features that are most contributing to prediction outcome and thus should be investigated for being causative variants for resistance mechanisms (Suppl. Table 3 & 4. First, with our ML approach we identified a list of common known resistance-associated mutations. For example, missense variants in the DNA gyrase gene *gyrA* are known to cause ofloxacin resistance, and mutations in the 16S ribosomal RNA gene *rrs* and small ribosomal subunit protein gene *rpsL* are causative for aminoglycosides resistance. Futhermore, mutations in the catalase-peroxidase gene *katG* that are known to cause resistance to isoniazid and prothionamide, and in the arabinosyltransferase A gene *embA* cause resistance to ethambutol. In several cases we observed SNPs upstream of known resistance-associated genes, which may signify the importance of regulation of gene expression in resistance, also reported in literature (36–38). In particular, we identified -15C>T (15 bases upstream of the start codon) mutation in the *fabG1* gene that is known to be associated with ethionamide resistance and -165T>C mutation in *rpsL* gene that is associated with aminoglycoside resistance. Moreover, we observed new connections of known resistance markers with other antibiotics. For example, *EmbA* is known to be associated with resistance to ethambutol (39), which is a first-line drug, but we see a Pro913Ser mutation to be predictive of resistance to ethionamide. Since ethionamide is a second-line drug, we can hypothesize that the corresponding strains are also resistant to ethambutol, indicating a case of multi-resistance, which is common for Mtb (40). In addition to known resistance-associated variants, we identified previously undescribed variants resulting in non-synonymous mutations: Ile145Met in a probable amino acid aminotransferase *PabC*, Val151Ala in lipoprotein *LppB*, Thr176Ile in a GntR family transcriptional regulator Rv3060c, Lys179Gln in a zinc metalloprotease *FtsH*, and Asn378Asp in a putative anion transporter ATPase Rv3679. To predict the impact of non-synonymous coding variants on the proteins’ function we analyzed their location in the protein three-dimensional structure with StructMAn (41). This tool predicts location of mutated position with respect to potential binding interfaces as well as protein core based on analysis of all experimentally resolved structures of complexes of homologous proteins. Thus, even if a certain binding event has never been detected for a particular bacterium, we can hypothesize about them by transferring information from related species. Most known resistance-associated mutations as well as previously undescribed variants can be mapped onto a structure where they lie on an interaction interface or in the protein core. For some new variants, there are neither any experimentally resolved structures of the corresponding protein, nor of its homologs. In this case we relied on structures predicted with AlphaFold, but four of six of these variants are classified as surface. No complexes of these proteins have been resolved, so it is possible that these surface mutations in fact take part in some biologically important interactions (Table 3).

**Table 3.** Structural classification of known resistance-associated and novel predictive variants.

| Variant ID | Gene | Mutation | Uniprot ID | RIN-based simple classification | Observed associated with resistance to | Reported to be associated with resistance to |
| --- | --- | --- | --- | --- | --- | --- |
| **Known resistance-associated mutations** |  |  |  |  |  |  |
| (7570, 'C,T', 'snp') | *GyrA* | A90V | P9WG47 | Ligand interaction | Ofloxacin, Ethionamide | Fluoroquinolone (42, 43) |
| (7362, 'G,C', 'snp') | *GyrA* | E21Q | P9WG47 | Protein interaction | Ofloxacin | Fluoroquinolone (42, 43) |
| (7585, 'G,C', 'snp') | *GyrA* | S95T | P9WG47 | DNA interaction | Ofloxacin | Fluoroquinolone (42, 43) |
| (781687, 'A,G', 'snp') | *RpsL* | K43R | P9WH63 | RNA interaction | Streptomycin | Streptomycin (42, 43) |
| (781822, 'A,G', 'snp') | *RpsL* | K88R | P9WH63 | RNA interaction | Streptomycin | Streptomycin (42, 43) |
| (2154724, 'C,A', 'snp') | *KatG* | R463L | P9WIE5 | Surface | Capreomycin, Ethionamide, Streptomycin | Isoniazid, Prothionamide (42, 43) |
| (4245969, 'C,T', 'snp') | *EmbA* | P913S | P9WNL9 | Core | Ethionamide | Ethambutol (42, 43) |
| **Previously unreported variants** |  |  |  |  |  |  |
| (906857, 'A,G', 'snp') | *PabC* | I145M | Q79FW0 | Protein interaction | Ethionamide, Ofloxacin | - |
| (4052349, 'T,G', 'snp') | *FtsH* | K179Q | P9WQN3 | Protein interaction | Ethionamide | - |
| (2867575, 'T,C', 'snp') | *LppB* | V151A | P9WK79 | Core | Ethionamide | - |
| **Previously unreported variants with no structural templates (AlphaFold(44) models used)** |  |  |  |  |  |  |
| (3422687, 'G,A', 'snp') | GntR family transcriptional regulator | T176I | P95098 | Core | Ethionamide | - |
| (4120926, 'A,G', 'snp') | anion transporter *ATPase* | N378D | I6Y498 | Surface | Ethionamide | - |
| (1896581, 'T,C', 'snp') | membrane protein | M36T | O53918 | Surface | Ethionamide | - |
| (623163, 'C,T', 'snp') | *PE_PGRS6* | A124V | L0T3X8 | Surface | Streptomycin | - |
| (623472, 'A,G', 'snp') | *PE_PGRS6* | D227G | L0T3X8 | Core | Streptomycin | - |
| (835611, 'C,T', 'snp') | hypothetical protein | T153M | I6X9N8 | Surface | Ofloxacin | - |

For example, mutation Ile145Met in the putative amino acid aminotransferase *PabC* (corresponds to SNP 906857A>G) is located on a protein-protein interaction interface of the protein (Fig 3a), as well as mutation Lys179Gln in the membrane-bound zinc metalloprotease *FtsH* (Fig 3b). Although the role of these proteins in ethionamide resistance is not clear, these mutations may change affinity of interaction between the subunits, and thus impact protein function. Moreover, *FtsH* has been linked to antibiotic resistance of other bacteria. In particular, mutations in *ftsH* and *ftsH*-controlling genes in *P. aeruginosa* increase sensitivity to tobramycin (45). In Staphylococcus aureus, *FtsH* is involved in bacterial virulence, stress resistance and *β*-lactam resistance (46). In *E. coli*, some isolates resistant to vancomycin and erythromycin also carry mutations in *ftsH* (47). The diversity of these compounds indicates that the mechanism of resistance in this case is probably related to membrane permeability.

**Fig. 3.**
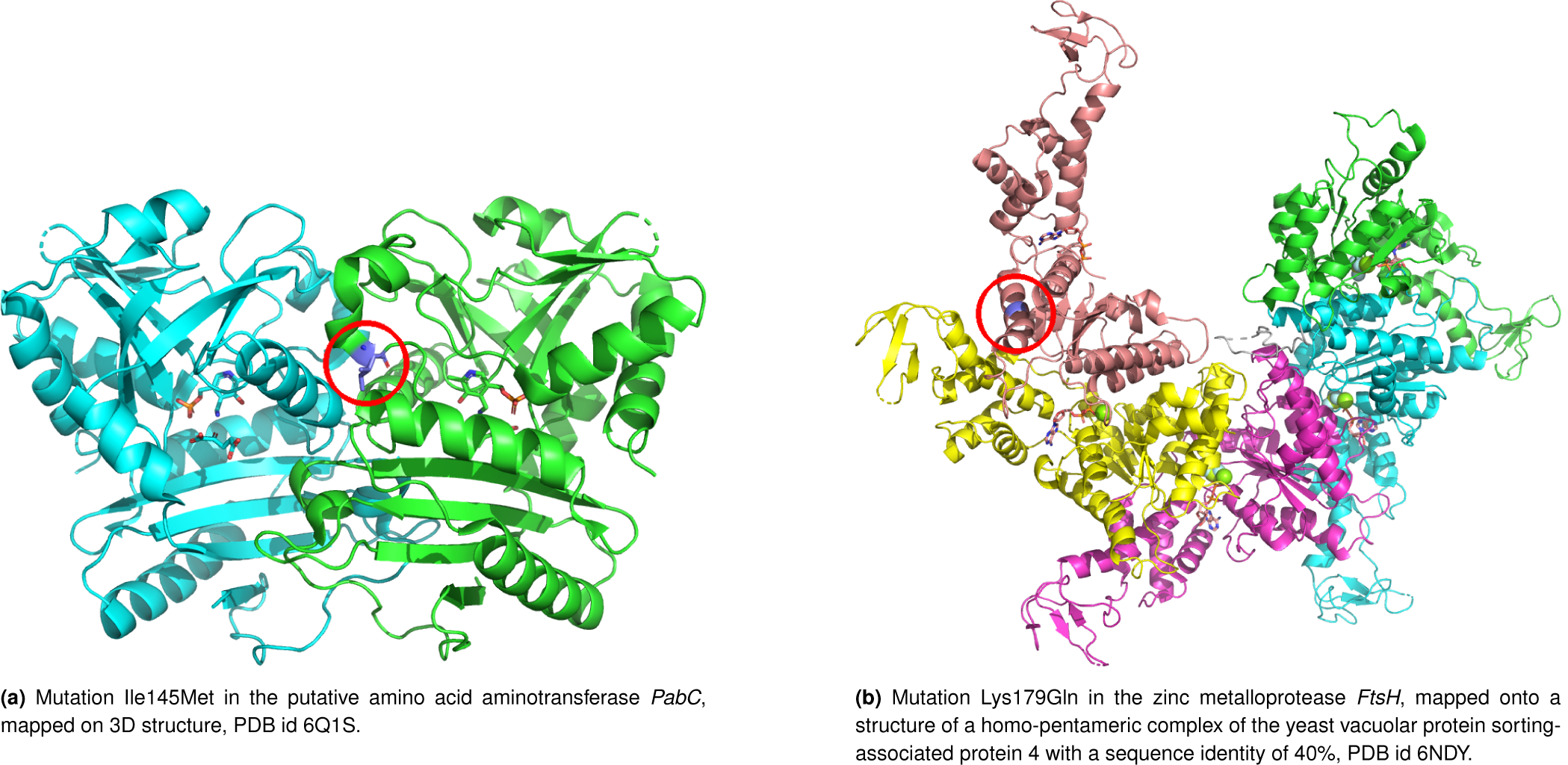
Structural analysis of the novel potential resistance markers. Corresponding mutations are shown as violet sticks, different chains in multimeric protein complexes are shown with ribbons in different colors. Figures were created with PyMOL (48)

## Discussion

Large collections of M. tuberculosis genomes provide a basis to study genomic features that cause antibiotic resistance in their populations. Understanding of molecular mechanisms, how both classical and new antitubercular drugs work and identifying specific mutations that allow Mtb to escape the effects of these drugs are important for development of new extended diagnostic panels as well as new efficient methods of treatments.

Traditional pipelines for Mtb drug resistance are based on calling known resistance-associated mutations in the newly sequenced genomic data (direct association), and work quite well with nearly perfect specificity. Meanwhile, ML approaches can improve this performance further (22). However, this study focussed only on 23 genes known to be associated with drug resistance, considering only on a small number of arguably most relevant features, and thus missing the possibility to discover novel resistance mechanisms. ML methods show promise not only for Mtb drug resistance, but for work on other pathogens (21), slowly making their way into personalized clinical guidelines (49).

In this work we propose a set of ML models for predicting antibiotics resistance in Mtb trained on public data. Despite the fact that our models are agnostic of prior biological knowledge on resistance markers and mechanisms, they were able to re-discover many of these known resistance markers. In addition to that, we have observed previously unreported resistance-associated mutations: in particular mutations in the membrane-bound metalloprotease *FtsH* associated with resistance to various antibiotics in various pathogens. This putative new genetic resistance marker in Mtb is likely associated with a new mechanism related to membrane permeation. In another instance we detected markers known to be associated with resistance to other antibiotics than those that were used in the phenotypic screens under consideration: for example, in the data for ethionamide (second-line drug) we detected a mutation in *EmbA* that is associated with resistance to a first-line drug ethambutol, which may hint at multi-resistance. This emphasizes the importance of using whole-genome sequences in such models as they are a mine of additional information.

Using whole-genome sequences, however, is connected with a danger of discovering many false-positive associations, due to low recombination rates and strong population structure in bacteria (11). Different approaches have been suggested to account for this: using linear mixed models (11), adding a feature that reflects predicted functional impact of the genetic variant (25), or training while accounting for population structure in the sample (21). We proposed a simple procedure and measure (parallelism-related phylogenetic score, PRPS) to rank genetic variants by their propensity to occur at multiple locations within the species’ phylogenetic tree and show that using only top-PRPS features improves performance of the models and minimizes the number of potential false positives. Combinations of multiple such scores are thinkable: for example, with predicted functional impact of corresponding mutations (50–52) or with their potential impact on protein structure and interactions (41).

Finally, ML methods are not restricted to SNPs. Whereas SNPs play a major role for resistance development in Mtb, other mechanisms are prevalent in other pathogens, such as horizontal gene transfer of resistance-associated genes via plasmids. Accordingly, features related to gene presence can be easily incorporated into models. Thanks to advances and increasing accessibility of long-read sequencing technologies, other potential mechanisms can be included and their impact on resistance can be explored: gene copy number, chromosomal rearrangements etc. (53–55). Whereas sequencing becomes cheaper, phenotypic screens that are required to train ML models are still laborious and costly, hence algorithmic developments to make most out of limited data are essential, too.

## Code

Implementation of the models and data tables can be found on GitHub: https://github.com/AlperYurtseven/ML-PRPS-MTB

## ACKNOWLEDGEMENTS

A.Y. and O.V.K. acknowledge financial support from the Klaus Faber Foundation.

A.A.A. was funded by the HelmholtzAI project AMR-XAI.

The work of O.O.B. is funded by Fonds zur Förderung der Wissenschaftlichen Forschung (FWF), Grant ESP 253-B.

