## Supplement for "Machine learning and phylogenetic analysis allow for predicting antibiotic resistance in *M. tuberculosis*"

**Table 1.** Genes associated with antibiotic resistance to different drugs from previous studies

| Drug name | Line of therapy | Pharmacological group | Gene name | References |
|---|---|---|---|---|
| Streptomycin | First line | Aminoglycosides | gidB, rrs, tlyA, rpsL | (41, 42) |
| Amikacin | Second line | Aminoglycosides | gidB, rrs, tlyA | (41, 42) |
| Capreomycin | Second line | Aminoglycosides | gidB, rrs, tlyA | (41, 42) |
| Kanamycin | Second line | Aminoglycosides | gidB, rrs, tlyA, eis | (41, 42) |
| Ofloxacin | Second line | Fluoroquinolones | gyrA, gyrB | (41, 42) |
| Ethionamide | Second line | Nicotinamide derivative | ethA, ethR, inhA, fabG1 | (41) |

**Table 2.** Significantly associated mutations found with pyseer.

| Mutation | LRT P-value | Product | Gene | Start-End |
|---|---|---|---|---|
| **Amikacin ((bonferroni corrected p-value threshold = 0.05/(no of variants tested) = 0.05/6551 = 7.63 x 10e-6))** | | | | |
| (1473246, 'A,G', 'snp')(A1401G) | 3.77E-91 | 16S ribosomal RNA | rrs | 1471845-1473382 |
| (1605149, 'C,T', 'snp')(P91L) | 6.94E-07 | hypothetical protein | - | 1604878-1606146 |
| **Capreomycin (bonferroni corrected p-value threshold = 0.05/(no of variants tested) = 0.05/6479 = 7.72 x 10e-6)** | | | | |
| (1473246, 'A,G', 'snp')(A1401G) | 5.35E-61 | 16S ribosomal RNA | rrs | 1471845-1473382 |
| **Ethionamide (bonferroni corrected p-value threshold = 0.05/(no of variants tested) = 0.05/6821 = 7.33 x 10e-6)** | | | | |
| (1673425, 'C,T', 'snp')(C-15T) | 3.29E-11 | Upstream of fabG1(1673440..1674183) | - | 1673440-1674183 |
| (1473246, 'A,G', 'snp')(A1401G) | 6.06E-07 | 16S ribosomal RNA | rrs | 1471845-1473382 |
| **Kanamycin (bonferroni corrected p-value threshold = 0.05/(no of variants tested) = 0.05/6505 = 7.69 x 10e-6)** | | | | |
| (1473246, 'A,G', 'snp')(A1401G) | 3.30E-51 | 16S ribosomal RNA | rrs | 1471845-1473382 |
| (2715346, 'G,A', 'snp')(C-14T) | 3.99E-13 | Downstream of eis(2714124..2715332) | - | 2714124-2715332 |
| (2715342, 'C,T', 'snp')(G-10A) | 1.02E-09 | Downstream of eis(2714124..2715332) | - | 2714124-2715332 |
| **Ofloxacin (bonferroni corrected p-value threshold = 0.05/(no of variants tested) = 0.05/7102 = 7.04 x 10e-6)** | | | | |
| (7585, 'G,C', 'snp')(S95T) | 2.02E-22 | DNA gyrase subunit A | gyrA | 7302-9818 |
| (7570, 'C,T', 'snp')(A90V) | 2.44E-18 | DNA gyrase subunit A | gyrA | 7302-9818 |
| **Streptomycin (bonferroni corrected p-value threshold = 0.05/(no of variants tested) = 0.05/7916 = 6.32 x 10e-6)** | | | | |
| (781687, 'A,G', 'snp')(K43R) | 8.43E-47 | 30S ribosomal protein S12 | rpsL | 781560-781934 |
| (781822, 'A,G', 'snp')(K88R) | 4.49E-30 | 30S ribosomal protein S12 | rpsL | 781560-781934 |
| (1472359, 'A,C', 'snp')(A514C) | 1.41E-22 | 16S ribosomal RNA | rrs | 1471845-1473382 |
| (2155168, 'C,G', 'snp')(S315T) | 1.41E-15 | catalase-peroxidase | katG | 2153889-2156111 |
| (2719057, 'G,A', 'snp')(G-26A) | 8.40E-07 | Upstream of hypothetical protein | - | 2719083-2719355 |
| (1673425, 'C,T', 'snp')(C-15T) | 9.24E-07 | Upstream of fabG1(1673440..1674183) | - | 1673440-1674183 |
| (4187063, 'G,A', 'snp')(G144R) | 1.97E-06 | AraC/XylS family transcriptional regulator | - | 4186634-4187695 |
| (1674048, 'G,A', 'snp')(L203L) | 3.50E-06 | 3-oxoacyl-ACP reductase FabG | fabG1 | 1673440-1674183 |

**Table 3.** Structural classification of known resistance-associated and novel predictive variants SVM.

| | Type | Effect | Gene | Product | Score ($r^2$) | CARD Database | Uniprot Accession |
|---|---|---|---|---|---|---|---|
| **Amikacin** | | | | | | | |
| (1472362, 'C,T', 'snp') | rRNA | non coding transcript variant | rrs | 16S ribosomal RNA | 0.187 +/- 0.000 | Aminoglycoside | - |
| **Capreomycin** | | | | | | | |
| (1472362, 'C,T', 'snp') | rRNA | non coding transcript variant | rrs | 16S ribosomal RNA | 0.169 +/- 0.000 | Aminoglycoside | - |
| **Ethionamide** | | | | | | | |
| (906857, 'A,G', 'snp') | CDS | missense variant c,435A>G p,Ile145Met | pabC | 4-amino-4-deoxychorismate lyase | 0.074 +/- 0.000 | - | Q79FW0 |
| (1472362, 'C,T', 'snp') | rRNA | non coding transcript variant | rrs | 16S ribosomal RNA | 0.069 +/- 0.000 | Aminoglycoside | - |
| (2867575, 'T,C', 'snp') | CDS | missense variant c,452T>C p,Val151Ala | lppB | lppB lipoprotein LppB | 0.048 +/- 0.000 | - | P9WK79 |
| (1673425, 'C,T', 'snp') | - | - | upstream of fabG1 | - | 0.048 +/- 0.000 | Ethionamide | P9WGT3 |
| (3422687, 'G,A', 'snp') | CDS | missense variant c,527C>T p,Thr176Ile | - | GntR family transcriptional regulator | 0.042 +/- 0.000 | - | P95098 |
| (4052349, 'T,G', 'snp') | CDS | missense variant c,535A>C p,Lys179Gln | ftsH | zinc metalloprotease FtsH | 0.037 +/- 0.000 | - | P9WQN3 |
| (4120926, 'A,G', 'snp') | CDS | missense variant c,1132A>G p,Asn378Asp | - | anion transporter ATPase | 0.037 +/- 0.000 | - | I6Y498 |
| (3732154, 'G,A', 'snp') | CDS | synonymous variant c,4782C>T p,Gly1594Gly | PPE54 | PPE family protein PPE54 | 0.037 +/- 0.000 | - | Q6MWY2 |
| **Kanamycin** | | | | | | | |
| (1472362, 'C,T', 'snp') | rRNA | non coding transcript variant | rrs | 16S ribosomal RNA | 0.147 +/- 0.000 | Aminoglycoside | - |
| (2165503, 'T,A', 'snp') | CDS | synonymous variant c,1809A>T p,Ala603Ala | PPE34 | PPE family protein PPE34 | 0.036 +/- 0.000 | - | Q79FI9 |
| **Ofloxacin** | | | | | | | |
| (7570, 'C,T', 'snp') | CDS | missense variant c,269C>T p,Ala90Val | gyrA | DNA gyrase subunit A | 0.125 +/- 0.000 | Fluoroquinolone | P9WG47 |
| (7362, 'G,C', 'snp') | CDS | missense variant c,61G>C p,Glu21Gln | gyrA | DNA gyrase subunit A | 0.076 +/- 0.000 | Fluoroquinolone | P9WG47 |
| (906857, 'A,G', 'snp') | CDS | missense variant c,435A>G p,Ile145Met | - | 4-amino-4-deoxychorismate lyase | 0.068 +/- 0.000 | - | Q79FW0 |
| (835611, 'C,T', 'snp') | CDS | missense variant c,458C>T p,Thr153Met | - | hypothetical protein | 0.049 +/- 0.000 | - | I6X9N8 |
| **Streptomycin** | | | | | | | |
| (623163, 'C,T', 'snp') | CDS | missense variant c,371C>T p,Ala124Val | PE PGRS6 | PE-PGRS family protein PE PGRS6 | 0.057 +/- 0.000 | - | L0T3X8 |
| (623472, 'A,G', 'snp') | CDS | missense variant c,680A>G p,Asp227Gly | PE PGRS6 | PE-PGRS family protein PE PGRS6 | 0.054 +/- 0.000 | - | L0T3X8 |
| (781395, 'T,C', 'snp') | - | - | 165bp upstream of rpsL | - | 0.045 +/- 0.000 | Streptomycin | P9WH63 |
| (2154724, 'C,A', 'snp') | CDS | missense variant c,1388G>T p,Arg463Leu | katG | catalase-peroxidase | 0.038 +/- 0.000 | Isoniazid & Prothionamide | P9WIE5 |

**Table 4.** Structural classification of known resistance-associated and novel predictive variants RF.

| | Type | Effect | Gene | Product | Score ($r^2$) | CARD Database | Uniprot Accession |
|---|---|---|---|---|---|---|---|
| | | | **Amikacin** | | | | |
| (1472362, 'C,T', 'snp') | rRNA | non coding transcript variant | rrs | 16S ribosomal RNA | 0.894 +/- 0.000 | Aminoglycoside | - |
| | | | **Capreomycin** | | | | |
| (1472362, 'C,T', 'snp') | rRNA | non coding transcript variant | rrs | 16S ribosomal RNA | 0.420 +/- 0.000 | Aminoglycoside | - |
| (2154724, 'C,A', 'snp') | CDS | missense variant c,1388G>T p,Arg463Leu | katG | catalase-peroxidase | 0.056 +/- 0.000 | Isoniazid & Prothionamide | P9WIE5 |
| | | | **Ethionamide** | | | | |
| (1670814, 'C,T', 'snp') | CDS | synonymous variant c,402C>T p,Gly134Gly | - | hypothetical protein | 0.130 +/- 0.000 | - | P9WLX5 |
| (1472362, 'C,T', 'snp') | rRNA | non coding transcript variant | rrs | 16S ribosomal RNA | 0.122 +/- 0.000 | Aminoglycoside | - |
| (7570, 'C,T', 'snp') | CDS | missense variant c,269C>T p,Ala90Val | gyrA | DNA gyrase subunit A | 0.107 +/- 0.000 | Fluoroquinolone | P9WG47 |
| (2154724, 'C,A', 'snp') | CDS | missense variant c,1388G>T p,Arg463Leu | katG | catalase-peroxidase | 0.045 +/- 0.000 | Isoniazid & Prothionamide | P9WIE5 |
| (4245969, 'C,T', 'snp') | CDS | missense variant c,2737C>T p,Pro913Ser | embA | arabinosyltransferase A | 0.042 +/- 0.000 | Ethambutol | P9WNL9 |
| (1722200, 'C,G', 'snp') | CDS | synonymous variant c,6210G>C p,Val2070Val | pks5 | polyketide synthase | 0.039 +/- 0.000 | - | O53901 |
| (1673425, 'C,T', 'snp') | - | - | upstream of fabG1 | - | 0.038 +/- 0.000 | Ethionamide | P9WGT3 |
| (1896581, 'T,C', 'snp') | CDS | missense variant c,107T>C p,Met36Thr | - | membrane protein | 0.036 +/- 0.000 | - | O53918 |
| | | | **Kanamycin** | | | | |
| (1472362, 'C,T', 'snp') | rRNA | non coding transcript variant | rrs | 16S ribosomal RNA | 0.722 +/- 0.000 | Aminoglycoside | - |
| | | | **Ofloxacin** | | | | |
| (7570, 'C,T', 'snp') | CDS | missense variant c,269C>T p,Ala90Val | gyrA | DNA gyrase subunit A | 0.464 +/- 0.000 | Fluoroquinolone | P9WG47 |
| (7362, 'G,C', 'snp') | CDS | missense variant c,61G>C p,Glu21Gln | gyrA | DNA gyrase subunit A | 0.451 +/- 0.000 | Fluoroquinolone | P9WG47 |
| (1983291, 'C,T', 'snp') | repeat region | synonymous variant c,1485G>A p,Pro495Pro | PPE24 | - | 0.035 +/- 0.000 | - | P9WI15 |
| | | | **Streptomycin** | | | | |
| (781395, 'T,C', 'snp') | - | - | 165bp upstream of rpsL | - | 0.255 +/- 0.000 | Streptomycin | P9WH63 |
| (2154724, 'C,A', 'snp') | CDS | missense variant c,1388G>T p,Arg463Leu | katG | catalase-peroxidase | 0.180 +/- 0.000 | Isoniazid & Prothionamide | P9WIE5 |
| (781687, 'A,G', 'snp') | CDS | missense variant c,128A>G p,Lys43Arg | rpsL | 30S ribosomal protein S12 | 0.055 +/- 0.000 | Streptomycin | P9WH63 |
| (1471659, 'C,T', 'snp') | ncRNA | non coding transcript variant | mcr3 | Putative small regulatory RNA | 0.045 +/- 0.000 | - | - |

**Table 5.** Faulty strains that are not used for this experiment

| | | | | | | |
|---|---|---|---|---|---|---|
| 1773.2540 | 1773.2720 | 1773.2950 | 1773.3400 | 1773.3490 | 1773.200 | 1773.2560 |
| 1773.2730 | 1773.2800 | 1773.2870 | 1773.2890 | 1773.3200 | 1773.3320 | 1773.3330 |
| 1773.3380 | 1773.3430 | 1773.3530 | 1773.3570 | 1773.3660 | 1773.370 | 1773.3770 |
| 1773.4990 | 1773.5000 | 1773.5010 | 1773.5020 | 1773.5030 | 1773.5040 | 1773.5050 |
| 1773.5070 | 1773.5080 | 1773.5090 | 1773.5100 | 1773.5120 | 1773.5130 | 1773.5140 |
| 1773.5160 | 1773.5170 | 1773.5180 | 1773.5190 | 1773.5200 | 1773.5210 | 1773.5220 |
| 1773.5240 | 1773.5250 | 1773.5260 | 1773.5280 | 1773.5290 | 1773.5300 | 1773.5310 |
| 1773.5330 | 1773.5340 | 1773.5350 | 1773.5360 | 1773.5370 | 1773.5380 | 1773.5390 |
| 1773.5410 | 1773.5420 | 1773.5430 | 1773.5440 | 1773.5450 | 1773.5930 | |

**Table 6.** Strains that are used for this experiment

| | | | | | | |
|---|---|---|---|---|---|---|
| 1423431.3 | 1773.15623 | 1773.5349 | 1773.3381 | 1773.4631 | 1448582.3 | 1773.14782 |
| 1448819.3 | 1773.4856 | 1773.4066 | 1295753.3 | 1773.1493 | 1773.2998 | 1773.2742 |
| 1773.5399 | 1773.15074 | 1773.15469 | 1773.2946 | 1773.255 | 1773.14764 | 1773.2858 |
| 1427196.3 | 1448512.3 | 1773.16089 | 1773.2944 | 1447451.3 | 1448730.3 | 1447516.3 |
| 1773.14979 | 1773.15457 | 1773.15812 | 1773.2786 | 1773.15727 | 1773.2562 | 1773.2836 |
| 1773.2735 | 1354165.3 | 1773.5725 | 1773.5573 | 1400873.3 | 1773.2947 | 1447520.3 |
| 1773.15906 | 1773.15534 | 1773.15155 | 1773.15142 | 1773.15687 | 1773.309 | 1773.5205 |
| 1773.15434 | 1773.5596 | 1423534.3 | 1773.5607 | 1402589.3 | 1773.5465 | 1448655.3 |
| 1773.2682 | 1354175.3 | 1773.5342 | 1773.4221 | 1438868.3 | 1773.3663 | 1773.14867 |
| 1773.14953 | 1773.15098 | 1773.15329 | 1422047.3 | 1773.422 | 1773.15086 | 1773.3391 |
| 1773.2721 | 1773.15579 | 1773.15788 | 1773.15291 | 1773.15905 | 1448404.3 | 1773.5249 |
| 1773.5351 | 1773.15823 | 1773.3179 | 1448443.3 | 1448462.3 | 1324276.3 | 1773.5366 |
| 1448648.3 | 1773.3516 | 1773.1476 | 1773.16088 | 1773.5268 | 1354106.3 | 1773.5145 |
| 1773.2986 | 1773.3832 | 1773.15369 | 1773.5184 | 1427292.3 | 1773.2539 | 1773.3526 |
| 1773.1508 | 1773.5568 | 1773.5663 | 1773.2461 | 1773.4211 | 1773.15701 | 1773.15836 |
| 1773.343 | 1773.3343 | 1773.15931 | 1773.15884 | 1773.3339 | 1773.15618 | 1397869.3 |
| 1773.5219 | 1455291.3 | 1773.15092 | 1773.591 | 1773.15595 | 1324252.3 | 1773.16026 |
| 1773.3392 | 1324224.3 | 1773.15974 | 1448662.3 | 1773.2708 | 1773.15219 | 1773.3385 |
| 1773.15935 | 1773.5545 | 1773.2768 | 1773.564 | 1773.1475 | 1448822.3 | 1773.15322 |
| 1773.15238 | 1448580.3 | 1423504.3 | 1773.15279 | 1423427.3 | 1447466.3 | 1773.15048 |
| 1295772.3 | 1773.5038 | 1267362.3 | 1448652.3 | 1773.15689 | 1448680.3 | 1773.3347 |
| 1447474.3 | 1324229.3 | 1773.5218 | 1408940.4 | 1773.15961 | 1448530.3 | 1427206.3 |
| 1448547.3 | 1773.16072 | 1773.5165 | 1397886.3 | 1427200.3 | 1773.15056 | 1773.15965 |
| 1773.2731 | 1448631.3 | 1773.4866 | 1773.3314 | 1397923.3 | 1408960.4 | 1773.3135 |
| 1773.15902 | 1773.572 | 1773.14998 | 1773.5467 | 1773.15295 | 1773.5659 | 1773.15281 |
| 1400925.3 | 1773.2759 | 1427210.3 | 1773.15787 | 1773.2799 | 1773.3486 | 1773.5425 |
| 1773.15143 | 1773.3216 | 1773.3155 | 1402584.3 | 1397905.3 | 1773.14738 | 1773.14954 |
| 1773.15801 | 1773.5579 | 1402600.3 | 1402592.3 | 1773.3007 | 1773.4995 | 1773.4996 |
| 1397938.3 | 1773.15675 | 1773.4027 | 1773.16044 | 1773.158 | 1773.5649 | 1773.15567 |
| 1773.14935 | 1773.14808 | 1773.14962 | 1773.5461 | 1773.2664 | 1448588.3 | 1448463.3 |
| 1773.2737 | 1773.3336 | 1773.2707 | 1422080.3 | 1773.5073 | 1773.5259 | 1448596.3 |
| 1773.15413 | 1773.15521 | 1448465.3 | 1773.3113 | 1773.16087 | 1354102.3 | 1773.3475 |
| 1448565.3 | 1773.15437 | 1773.2471 | 1773.5681 | 1773.1533 | 1773.15793 | 1773.16018 |
| 1295795.3 | 1397913.3 | 1773.3358 | 1448427.3 | 1773.14947 | 1422040.3 | 1773.5561 |
| 1773.3138 | 1773.14916 | 1773.15171 | 1773.5043 | 1773.5718 | 1773.3042 | 1773.15755 |
| 1773.5035 | 1402583.3 | 1773.3459 | 1773.3163 | 1773.14754 | 1773.2671 | 1773.2781 |
| 1267357.3 | 1773.15706 | 1773.15033 | 1773.14876 | 1773.5237 | 1773.15769 | 1773.2872 |
| 1773.3227 | 1773.5186 | 1773.2969 | 1455283.3 | 1267358.3 | 1773.4851 | 1773.3268 |
| 1773.5884 | 1773.2848 | 1324219.3 | 1447447.3 | 1448610.3 | 1773.14889 | 1773.14835 |
| 1773.3164 | 1773.15735 | 1773.563 | 1773.4202 | 1267355.3 | 1397881.3 | 1408907.4 |
| 1397935.3 | 1773.2815 | 1773.5577 | 1773.2655 | 1354114.3 | 1773.5614 | 1773.5212 |
| 1773.15373 | 1773.15639 | 1773.2772 | 1447513.3 | 1448470.3 | 1423523.3 | 1402599.3 |
| 1400898.3 | 1354172.3 | 1773.2744 | 1773.5002 | 1773.15899 | 1773.5441 | 1773.15065 |
| 1773.15976 | 1773.2722 | 1427274.3 | 1773.15293 | 1773.5123 | 1773.5908 | 1773.1516 |
| 1773.15851 | 652616.4 | 1773.15506 | 1773.342 | 1423549.3 | 1773.1535 | 1448516.3 |
| 1773.1528 | 1773.3462 | 1773.2891 | 1773.16075 | 1773.3369 | 1773.2938 | 1773.3028 |
| 1773.5174 | 1773.2908 | 1773.5554 | 1773.5108 | 1773.15023 | 1773.3105 | 1773.15568 |
| 1773.3307 | 1773.2616 | 1773.3037 | 1773.3186 | 1773.346 | 1448687.3 | 1773.589 |
| 1773.2802 | 1773.28 | 1773.15085 | 1423499.3 | 1422038.3 | 1773.15471 | 1773.332 |
| 1773.5541 | 1773.3025 | 1324243.3 | 1448711.3 | 1773.2979 | 1773.4644 | 1773.266 |
| 1448810.3 | 1354126.3 | 1773.4223 | 1773.5583 | 1773.15504 | 1773.15345 | 1421993.3 |
| 1427304.3 | 1773.5201 | 1773.5893 | 1773.5481 | 1295798.3 | 1773.5408 | 1773.2904 |

**Table 6.** Strains that are used for this experiment

| | | | | | | |
|---|---|---|---|---|---|---|
| 1773.2658 | 1773.5152 | 1427216.3 | 1773.506 | 1400935.3 | 1773.2514 | 1773.5076 |
| 1773.15201 | 1773.322 | 1773.15632 | 1773.14873 | 1773.4255 | 1773.15942 | 1448630.3 |
| 1773.2876 | 1773.3253 | 1773.1481 | 1427310.3 | 1773.2753 | 1295751.3 | 1773.5292 |
| 1773.2985 | 1773.5348 | 1773.15859 | 1427262.3 | 1773.15308 | 1773.14736 | 1773.3271 |
| 1773.1487 | 1773.267 | 1773.423 | 1448455.3 | 1773.4261 | 1773.5548 | 1773.3308 |
| 1448805.3 | 1773.15726 | 1773.16081 | 1773.286 | 1773.15784 | 1448417.3 | 1448704.3 |
| 1773.5341 | 1773.5175 | 1324270.3 | 1773.15165 | 1295763.3 | 1773.5046 | 1773.2948 |
| 1773.2959 | 1773.15371 | 1423502.3 | 1773.3353 | 1773.5322 | 1773.5538 | 1773.3177 |
| 1354111.3 | 1448612.3 | 1773.5198 | 1773.5615 | 1448722.3 | 1773.16102 | 1773.14863 |
| 1773.5293 | 1418256.3 | 1400879.3 | 1773.15362 | 1773.16096 | 1773.15853 | 1422016.3 |
| 1324271.3 | 1773.529 | 1773.15821 | 1773.3324 | 1773.3187 | 1397929.3 | 1773.1512 |
| 1773.157 | 1773.2835 | 1773.334 | 1397855.3 | 1773.5319 | 1448651.3 | 1773.4616 |
| 1773.15739 | 1400884.3 | 1773.16019 | 1773.14922 | 1773.3383 | 1773.1595 | 1773.537 |
| 1773.3348 | 1773.5295 | 1773.15709 | 1324290.3 | 1773.15712 | 1773.3356 | 1773.15594 |
| 1773.3404 | 1773.15745 | 1773.3685 | 1354127.3 | 1773.3313 | 1400909.3 | 1773.14739 |
| 1354135.3 | 1773.15359 | 1773.306 | 1773.4164 | 1773.16048 | 1773.2639 | 1773.15138 |
| 1773.296 | 1773.15344 | 1773.15224 | 1773.317 | 1773.2559 | 1773.2967 | 1773.5122 |
| 1773.16028 | 1773.2907 | 1422084.3 | 1295740.3 | 1447501.3 | 1427330.3 | 1773.5515 |
| 1773.15062 | 1773.2436 | 1773.15667 | 1773.519 | 1773.15686 | 1448633.3 | 1773.5118 |
| 1423514.3 | 1773.15981 | 1354137.3 | 1773.2494 | 1423510.3 | 1773.2608 | 1448634.3 |
| 1773.3597 | 1773.1537 | 1427261.3 | 1773.5163 | 1448783.3 | 1773.15167 | 1773.15338 |
| 1773.15862 | 1773.14796 | 1773.3512 | 1427325.3 | 1773.275 | 1773.546 | 1427322.3 |
| 1408932.4 | 33894.5 | 1773.15406 | 1448394.3 | 1773.16002 | 1773.3421 | 1773.16043 |
| 1448697.3 | 1773.347 | 1773.2521 | 1773.4219 | 1773.1536 | 1773.14855 | 1773.3092 |
| 1423506.3 | 1423539.3 | 1773.499 | 1447449.3 | 1447488.3 | 1773.14886 | 1773.5331 |
| 1354132.3 | 1773.334 | 1423425.3 | 1773.5095 | 1448738.3 | 1773.2893 | 1773.3087 |
| 1773.2554 | 1422100.3 | 1773.15251 | 1773.2874 | 1773.3053 | 1773.515 | 1773.14853 |
| 1773.5507 | 1773.3346 | 1773.4855 | 1773.2612 | 1773.15585 | 1397863.3 | 1773.3444 |
| 1773.5458 | 1773.4208 | 1773.3522 | 1408974.4 | 1423520.3 | 1773.2489 | 1773.5139 |
| 1773.4321 | 1773.5657 | 1422029.3 | 1773.3282 | 1773.15464 | 1773.289 | 1448617.3 |
| 1773.2706 | 1400870.3 | 1773.2439 | 1773.5469 | 1773.3056 | 1773.5707 | 1773.15218 |
| 1448645.3 | 1438837.3 | 1438839.3 | 1324235.3 | 1773.5624 | 1773.1486 | 1773.2649 |
| 1773.15877 | 1397861.3 | 1448597.3 | 1773.15263 | 1448641.3 | 1773.15375 | 1773.5365 |
| 1448732.3 | 1773.15454 | 1773.2541 | 1773.2469 | 1773.5921 | 1324225.3 | 1773.1573 |
| 1448795.3 | 1448694.3 | 1773.2523 | 1773.4989 | 1773.3491 | 1448791.3 | 1773.5771 |
| 1773.328 | 1773.2971 | 1773.5414 | 1773.15777 | 1773.3172 | 1773.5453 | 1773.2784 |
| 1448627.3 | 1773.15144 | 1448787.3 | 1773.3036 | 1773.324 | 1448757.3 | 1773.14818 |
| 1773.15229 | 1773.4535 | 1773.5162 | 1447509.3 | 1773.15031 | 1773.15614 | 1773.3088 |
| 1773.15447 | 1773.333 | 1773.15636 | 1773.14903 | 1773.3406 | 1773.15805 | 1408936.4 |
| 1773.15284 | 1773.15306 | 1773.16013 | 1408965.4 | 1448827.3 | 1773.14806 | 1773.5379 |
| 1773.2779 | 1447496.3 | 1400933.3 | 1295792.3 | 1427252.3 | 1773.15876 | 1773.15882 |
| 1773.245 | 1773.5906 | 1773.2796 | 1773.2582 | 1773.16016 | 1773.3015 | 1427281.3 |
| 1448473.3 | 1773.3267 | 1773.5387 | 1773.5261 | 1448399.3 | 1354101.3 | 1448671.3 |
| 1773.15937 | 1295726.3 | 1773.15729 | 1773.2983 | 1773.5604 | 1773.3396 | 1773.15005 |
| 1773.5179 | 1773.5555 | 1773.2468 | 1448778.3 | 1448430.3 | 1773.5364 | 1773.3467 |
| 1773.15828 | 1438849.3 | 1773.2879 | 1773.14931 | 1447435.3 | 1773.5109 | 1773.14958 |
| 1773.16074 | 1448406.3 | 1773.5407 | 1397860.3 | 1773.5224 | 1773.2776 | 1773.3182 |
| 1773.3153 | 1427319.3 | 1773.15184 | 1773.15079 | 1773.501 | 1448586.3 | 1773.14978 |
| 1773.2622 | 1773.3259 | 1773.1494 | 1773.14951 | 1773.5182 | 1773.14898 | 1773.2524 |
| 1324264.3 | 1773.326 | 1773.15282 | 1417011.3 | 1773.338 | 1773.2766 | 1773.4248 |
| 1448567.3 | 1447436.3 | 1773.3694 | 1773.15888 | 1773.3258 | 1773.15334 | 1773.3364 |
| 1438863.3 | 1773.15517 | 1408937.4 | 1773.3074 | 1773.5029 | 1773.2578 | 1773.3408 |

**Table 6.** Strains that are used for this experiment

| | | | | | | |
|---|---|---|---|---|---|---|
| 1773.5719 | 1773.15361 | 1427312.3 | 1417018.3 | 1400940.3 | 1438843.3 | 1448640.3 |
| 1400930.3 | 1354155.3 | 1773.503 | 1773.3454 | 1354120.3 | 1773.3114 | 1773.3251 |
| 1400871.3 | 1773.3357 | 1773.5333 | 1773.4228 | 1773.15772 | 1773.59 | 1773.3151 |
| 1773.2634 | 1773.2989 | 1773.363 | 1773.43 | 1773.15479 | 1448632.3 | 1773.15298 |
| 1773.5521 | 1448434.3 | 1773.5433 | 1773.2982 | 1773.15059 | 1773.5164 | 1773.3688 |
| 1773.2853 | 1773.14974 | 1773.4667 | 1773.5083 | 1773.5241 | 1773.16038 | 1773.5648 |
| 1773.296 | 1400886.3 | 1773.15699 | 1773.5383 | 1773.5508 | 1427307.3 | 1773.1491 |
| 1773.4658 | 1773.15159 | 1773.2643 | 1773.14755 | 1773.3333 | 1448491.3 | 1773.15057 |
| 1773.3703 | 1773.15814 | 1773.2691 | 1773.329 | 1773.5647 | 1417009.3 | 1448821.3 |
| 1267360.3 | 1422079.3 | 1455266.3 | 1397901.3 | 1773.5567 | 1397897.3 | 1773.3249 |
| 1427209.3 | 1773.14983 | 1773.5172 | 1427250.3 | 1773.15575 | 1773.2566 | 1773.2673 |
| 1408927.4 | 1773.15909 | 1773.2618 | 1427207.3 | 1773.15496 | 1423554.3 | 1773.5033 |
| 1400874.3 | 1773.5042 | 1773.15197 | 1773.15307 | 1773.16069 | 1773.16077 | 1773.2477 |
| 1773.31 | 1773.4031 | 1438881.3 | 1773.3418 | 1455263.3 | 1773.15102 | 1773.199 |
| 1773.312 | 1773.3402 | 1438842.3 | 1773.15529 | 1773.15679 | 1354116.3 | 1773.5302 |
| 1773.2683 | 1773.5142 | 1421997.3 | 1773.2736 | 1354107.3 | 1773.4236 | 1400921.3 |
| 1773.14915 | 1773.4241 | 1773.2647 | 1455278.3 | 1324277.3 | 1773.333 | 1773.5602 |
| 1773.2995 | 1773.2486 | 1773.14893 | 1773.2642 | 1423516.3 | 1773.3184 | 1773.3201 |
| 1773.3012 | 1773.5084 | 1773.2987 | 1423428.3 | 1773.15409 | 1773.15682 | 1408918.4 |
| 1400924.3 | 1773.2728 | 1773.5251 | 1773.15638 | 1773.5313 | 1773.15356 | 1773.5041 |
| 1448807.3 | 1773.2687 | 1773.14762 | 1773.34 | 1773.2452 | 1773.15954 | 1773.285 |
| 1773.15725 | 1773.5713 | 1421966.3 | 1773.486 | 1773.3142 | 1773.3171 | 1773.4671 |
| 1427221.3 | 1773.5473 | 1427286.3 | 1773.15052 | 1773.2764 | 1773.2831 | 1773.15016 |
| 1773.2747 | 1773.14913 | 1773.5459 | 1773.5697 | 1773.15527 | 1773.3102 | 1773.2718 |
| 1455311.3 | 1773.3131 | 1773.15025 | 1773.15874 | 1773.15047 | 1447444.3 | 1422069.3 |
| 1773.15841 | 1773.2725 | 1773.3513 | 1408967.4 | 1773.516 | 1773.5355 | 1773.151 |
| 1773.5885 | 1773.5239 | 1455308.3 | 1773.14929 | 1773.15207 | 1773.5101 | 1773.14791 |
| 1402595.3 | 1773.4598 | 1773.251 | 1773.3132 | 1773.2481 | 1773.3805 | 1773.15538 |
| 1773.15401 | 1773.2656 | 1773.5724 | 1773.3432 | 1324228.3 | 1773.5194 | 1773.15995 |
| 1773.14793 | 1354119.3 | 1773.3523 | 1773.5024 | 1427219.3 | 1773.2873 | 1417029.3 |
| 1773.5196 | 1773.15939 | 1773.557 | 1773.5327 | 1773.2446 | 1773.3683 | 1773.15957 |
| 1773.3248 | 1773.5317 | 1773.3424 | 1448690.3 | 1773.5661 | 1773.14923 | 1455292.3 |
| 1422098.3 | 1773.2677 | 1447465.3 | 1773.5204 | 1773.15254 | 1427257.3 | 1773.15318 |
| 1773.3305 | 1773.3162 | 1448439.3 | 1773.2637 | 1773.3083 | 1773.3754 | 1354123.3 |
| 1397878.3 | 1448615.3 | 1773.15449 | 1773.244 | 1448428.3 | 1773.15809 | 1773.2662 |
| 1773.15549 | 1773.3269 | 1773.5535 | 1773.15768 | 1773.4225 | 1773.2589 | 1408963.4 |
| 1423573.3 | 1448743.3 | 1773.15163 | 1397880.3 | 1773.1507 | 1773.15843 | 1773.4676 |
| 1773.2571 | 1773.15834 | 1773.15 | 1773.331 | 1773.2698 | 1773.5501 | 1773.14992 |
| 1773.15536 | 1773.2757 | 1324232.3 | 1448616.3 | 1448643.3 | 1773.2867 | 1448623.3 |
| 1773.4026 | 1773.2743 | 1773.2719 | 1448480.3 | 1773.538 | 1773.343 | 1773.5493 |
| 1773.15332 | 1773.32 | 1408928.4 | 1773.4099 | 1773.3206 | 1773.2638 | 1773.5622 |
| 1427249.3 | 1773.15578 | 1773.15858 | 1773.1506 | 1427227.3 | 1448663.3 | 1773.5087 |
| 1423483.3 | 1773.5621 | 1324250.3 | 1295747.3 | 1773.16061 | 1773.15475 | 1773.5353 |
| 1773.26 | 1773.2762 | 1773.2838 | 1397890.3 | 1448808.3 | 1448482.3 | 1773.15278 |
| 1773.16032 | 1773.1498 | 1448577.3 | 1773.15598 | 1773.2565 | 1773.34 | 1421937.3 |
| 1295721.3 | 1773.14819 | 1408946.4 | 1773.15091 | 1773.15237 | 1773.15685 | 1773.2792 |
| 1438865.3 | 1448744.3 | 1773.3502 | 1773.15762 | 1773.3051 | 1447457.3 | 1324288.3 |
| 1773.3442 | 1427289.3 | 1773.16036 | 1295782.3 | 1427246.3 | 1773.15582 | 1773.5345 |
| 1773.15695 | 1773.15456 | 1773.276 | 1773.1565 | 1773.5308 | 1773.15017 | 1773.3458 |
| 1423451.3 | 1773.5055 | 1773.16 | 1773.15615 | 1773.15069 | 1773.1532 | 1773.3062 |
| 1773.3411 | 1773.5323 | 1773.5682 | 1773.3104 | 1773.15034 | 1773.3974 | 1773.2845 |
| 1773.2604 | 1773.5729 | 1773.15616 | 1773.14996 | 1773.5901 | 1448800.3 | 1773.2599 |

**Table 6.** Strains that are used for this experiment

| | | | | | | |
|---|---|---|---|---|---|---|
| 1773.3024 | 1773.15134 | 1423437.3 | 1773.5914 | 1773.5173 | 1448431.3 | 1773.3241 |
| 1773.15013 | 1773.14804 | 1773.14817 | 1773.281 | 1408916.4 | 1773.3011 | 1773.408 |
| 1773.15335 | 1773.204 | 1773.2507 | 1773.15014 | 1773.5114 | 1447434.3 | 1773.4994 |
| 1773.5311 | 1423541.3 | 1773.15001 | 1773.15273 | 1397941.3 | 1773.1477 | 1773.2928 |
| 1448620.3 | 1773.535 | 1773.15393 | 1773.14881 | 1773.15407 | 1773.3158 | 1773.16066 |
| 1773.2923 | 1448529.3 | 1773.5257 | 1773.5601 | 1773.2528 | 1773.15436 | 1295756.3 |
| 1773.2579 | 1397909.3 | 1773.3449 | 1773.15794 | 1773.31 | 1773.5575 | 1773.3303 |
| 1773.15977 | 1773.293 | 1397916.3 | 1773.5557 | 1427235.3 | 1408924.4 | 1773.3359 |
| 1773.5085 | 1773.14899 | 1773.15721 | 1773.2789 | 1447476.3 | 1773.3335 | 1324260.3 |
| 1427189.3 | 1448678.3 | 1773.15495 | 1773.15389 | 1423481.3 | 1773.15959 | 1773.15766 |
| 1773.14949 | 1773.15055 | 1773.3316 | 1354145.3 | 1417010.3 | 1408955.4 | 1402590.3 |
| 1773.16054 | 1773.3471 | 1448668.3 | 1773.1521 | 1773.4246 | 1773.5675 | 1427323.3 |
| 1773.3372 | 1773.5384 | 1400937.3 | 1448669.3 | 1773.3243 | 1773.15039 | 1773.5484 |
| 1422012.3 | 1773.15924 | 1773.15374 | 1773.3289 | 1324275.3 | 1354191.3 | 1773.5026 |
| 1773.15952 | 1773.5017 | 1773.2503 | 1324237.3 | 1773.351 | 1448532.3 | 1295799.3 |
| 1773.534 | 1448407.3 | 1422036.3 | 1427268.3 | 1354143.3 | 1773.5032 | 1422021.3 |
| 1773.1578 | 1773.5576 | 1448672.3 | 1773.3079 | 1773.4209 | 1773.4865 | 1773.5556 |
| 1773.2459 | 1773.2964 | 1324281.3 | 1773.2869 | 1773.1583 | 1773.4992 | 1773.5479 |
| 1773.5421 | 1427202.3 | 1773.2686 | 1447504.3 | 1773.15182 | 1773.3531 | 1773.2517 |
| 1448595.3 | 1773.14742 | 1773.15663 | 1773.313 | 1773.15603 | 1773.3439 | 1773.3119 |
| 1773.3254 | 1427236.3 | 1773.5539 | 1421941.3 | 1773.249 | 1773.3436 | 1773.5111 |
| 1773.15149 | 1354105.3 | 1773.3069 | 1773.15833 | 1773.1545 | 1773.359 | 1773.3043 |
| 1354156.3 | 1423461.3 | 1773.16071 | 1354162.3 | 1448673.3 | 1773.14956 | 1773.15722 |
| 1773.5405 | 1423438.3 | 1427315.3 | 1773.5022 | 1773.349 | 1773.523 | 1773.15708 |
| 1397910.3 | 1324234.4 | 1773.14975 | 1773.3334 | 1773.5668 | 1773.14779 | 1773.15605 |
| 1324220.3 | 1421988.3 | 1422093.3 | 1438871.3 | 1448642.3 | 1408964.4 | 1773.5018 |
| 1438869.3 | 1402597.3 | 1324300.3 | 1773.14973 | 1773.15566 | 1773.15928 | 1773.3484 |
| 1397853.3 | 1773.5432 | 1324242.3 | 1427224.3 | 1773.5171 | 1324253.3 | 1773.5234 |
| 1773.327 | 1773.2601 | 1773.15106 | 1773.15491 | 1773.15478 | 1773.15494 | 1773.15811 |
| 1773.5157 | 1773.2652 | 1773.15419 | 1408970.4 | 1773.15114 | 1421996.3 | 1773.5028 |
| 1773.525 | 1448752.3 | 1773.2714 | 1438833.3 | 1773.15966 | 1773.1594 | 1427187.3 |
| 1773.3696 | 1773.4862 | 1773.5392 | 1773.3678 | 1773.5316 | 1773.2657 | 1773.1543 |
| 1773.3337 | 1773.3166 | 1773.2788 | 1448784.3 | 1773.2949 | 1773.15511 | 1397877.3 |
| 1773.15804 | 1773.15872 | 1773.2861 | 1773.5662 | 1397900.3 | 1773.4148 | 1423536.3 |
| 1773.15412 | 1354187.3 | 1773.3503 | 1773.256 | 1773.15629 | 1773.5066 | 1447502.3 |
| 1773.3077 | 1773.14926 | 1773.1538 | 1773.4104 | 1773.1534 | 1773.15822 | 1773.15038 |
| 1773.3524 | 1773.15022 | 1448734.3 | 1773.5471 | 1773.3045 | 1422061.3 | 1773.1599 |
| 1773.15938 | 1773.355 | 1455285.3 | 1448695.3 | 1397926.3 | 1448709.3 | 1773.16034 |
| 1773.3447 | 1773.1577 | 1773.15271 | 1773.15141 | 1773.2457 | 1773.2822 | 1448799.3 |
| 1773.14985 | 1773.1589 | 1773.325 | 1427278.3 | 1773.2775 | 1773.3021 | 1773.53 |
| 1773.5426 | 1773.15315 | 1773.14937 | 1773.5135 | 1447459.3 | 1773.2826 | 1448790.3 |
| 1400923.3 | 1397933.3 | 1423535.3 | 1423484.3 | 1773.15379 | 1773.3199 | 1448755.3 |
| 1773.2513 | 1448562.3 | 1773.15657 | 1773.15045 | 1773.15168 | 1773.2999 | 1447485.3 |
| 1773.14984 | 1773.5089 | 1773.4229 | 1773.253 | 1448600.3 | 1427179.3 | 1423530.3 |
| 1397893.3 | 1773.15274 | 1397924.3 | 1773.2827 | 1773.15577 | 1773.14737 | 1773.5256 |
| 1773.16004 | 1773.5529 | 1773.3493 | 1455293.3 | 1354153.3 | 1773.15199 | 1773.2623 |
| 1773.5448 | 1773.15357 | 1773.3157 | 1773.16029 | 1773.1479 | 1773.14829 | 1448552.3 |
| 1448637.7 | 1773.593 | 1773.2635 | 1773.2996 | 1773.3485 | 1773.2881 | 1773.15265 |
| 1773.3147 | 1773.5007 | 1448592.3 | 1773.2791 | 1773.4845 | 1773.5312 | 1773.15664 |
| 1773.2555 | 1397934.3 | 1773.15312 | 1773.15123 | 1773.14797 | 1773.323 | 1773.5635 |
| 1773.15223 | 1773.3223 | 1773.2479 | 1773.308 | 1773.2614 | 1773.5008 | 1448660.3 |
| 1773.14758 | 1773.518 | 1773.3103 | 1427225.3 | 1773.15641 | 1423555.3 | 1773.1567 |

**Table 6.** Strains that are used for this experiment

| | | | | | | |
|---|---|---|---|---|---|---|
| 1400936.3 | 1397872.3 | 1773.275 | 1773.15723 | 1773.14977 | 1773.5702 | 1773.4691 |
| 1773.2512 | 1773.5283 | 1773.4858 | 1773.2505 | 1773.3287 | 1773.16094 | 1773.315 |
| 1773.2941 | 1773.15544 | 1773.3455 | 1773.1605 | 1773.15376 | 1397857.3 | 1295801.3 |
| 1402601.3 | 1773.3302 | 1773.5487 | 1773.5148 | 1448740.3 | 1438860.3 | 1773.15651 |
| 1773.15994 | 1354152.3 | 1773.15126 | 1773.15767 | 1773.4207 | 1773.1557 | 1773.2777 |
| 1427314.3 | 1773.357 | 1773.5126 | 1773.5156 | 1773.5913 | 1773.5206 | 1773.5564 |
| 1773.5715 | 1423432.3 | 1354121.3 | 1773.3244 | 1773.14787 | 1773.4049 | 1773.3174 |
| 1773.14816 | 1773.15445 | 1773.2485 | 1773.532 | 1295803.3 | 1773.15466 | 1773.284 |
| 1773.5009 | 1773.2825 | 1773.2897 | 1773.5887 | 1773.14775 | 1408909.4 | 1773.16053 |
| 1773.15314 | 1438854.3 | 1422034.3 | 1773.15222 | 1773.3274 | 1773.3415 | 1773.5039 |
| 1773.14909 | 1773.15531 | 1448585.3 | 1773.14823 | 1773.14842 | 1773.15856 | 1773.15267 |
| 1448572.3 | 1448479.3 | 1354174.3 | 1773.2899 | 1773.5571 | 1773.5199 | 1397882.3 |
| 1773.15264 | 1773.5131 | 1397895.3 | 1773.15619 | 1773.3169 | 1773.2733 | 1354142.3 |
| 1773.15428 | 1422001.3 | 1773.14892 | 1773.14788 | 1447521.3 | 1773.2627 | 1773.14765 |
| 1773.2669 | 1773.5488 | 1773.15826 | 1773.3231 | 1773.15127 | 1773.2894 | 1438840.3 |
| 1773.15532 | 1773.15396 | 1400920.3 | 1773.15489 | 1324282.3 | 1773.14884 | 1773.5346 |
| 1448727.3 | 1773.15848 | 1448446.3 | 1773.2467 | 1773.2843 | 1427256.3 | 1421956.3 |
| 1773.377 | 1448703.3 | 1773.2833 | 1773.15626 | 1448767.3 | 1773.15728 | 1773.5422 |
| 1773.15989 | 1773.15019 | 1773.16091 | 1773.2864 | 1773.14843 | 1427272.3 | 1447452.3 |
| 1447454.3 | 1773.2535 | 1773.5047 | 1773.15512 | 1773.2586 | 1773.2756 | 1773.5578 |
| 1397915.3 | 1773.5656 | 1773.15969 | 1773.14826 | 1773.16006 | 1773.4991 | 1448656.3 |
| 1773.3136 | 1447493.3 | 1773.556 | 1427313.3 | 1773.1581 | 1397889.3 | 1448590.3 |
| 1773.292 | 1773.5644 | 1773.15658 | 1773.1513 | 1773.2727 | 1773.325 | 1773.3544 |
| 1354147.3 | 1773.15081 | 1773.5613 | 1400931.3 | 1773.14838 | 1773.5093 | 1324254.3 |
| 1773.15798 | 1773.3351 | 1773.14813 | 1455284.3 | 1423547.3 | 1448624.3 | 1773.4184 |
| 1773.2745 | 1447503.3 | 1421977.3 | 1448771.3 | 1267356.3 | 1773.2854 | 1773.15026 |
| 1773.2975 | 1423572.3 | 1773.14802 | 1773.2771 | 1773.15714 | 1448716.3 | 1773.5258 |
| 1773.15786 | 1448676.3 | 1773.313 | 1773.15569 | 1773.15733 | 1773.15443 | 1422022.3 |
| 1408954.4 | 1773.2787 | 1397864.3 | 1773.4214 | 1773.4252 | 1773.3483 | 1773.1553 |
| 1400929.3 | 1773.4993 | 1773.5217 | 1773.15718 | 1773.15508 | 1773.14894 | 1448537.3 |
| 1408923.4 | 1438832.3 | 1773.3842 | 1773.15277 | 1773.265 | 1773.1566 | 1448400.3 |
| 1402596.3 | 1400938.3 | 1773.2818 | 1773.15209 | 1773.3414 | 1324255.3 | 1773.5226 |
| 1773.36 | 1773.5606 | 1773.2451 | 1773.5496 | 1773.5284 | 1448691.3 | 1773.14921 |
| 1417027.3 | 1773.15964 | 1324263.3 | 1448628.3 | 1773.4201 | 1448677.3 | 1773.514 |
| 1773.15477 | 1773.261 | 1773.14732 | 1773.2924 | 1773.15458 | 1448598.3 | 1423430.3 |
| 1773.337 | 1427329.3 | 1408943.4 | 1773.14885 | 1773.5666 | 1773.5291 | 1427203.3 |
| 1773.14801 | 1448509.3 | 1773.2935 | 1773.14887 | 1773.1572 | 1773.3121 | 1773.2619 |
| 1423542.3 | 1438867.3 | 1773.3246 | 1773.5192 | 1423519.3 | 1773.2763 | 1427230.3 |
| 1773.3255 | 1447489.3 | 1773.4707 | 1773.5034 | 1448397.3 | 1773.14828 | 1773.559 |
| 1773.349 | 1397918.3 | 1773.15455 | 1427290.3 | 1773.3413 | 1773.15011 | 1773.14777 |
| 1773.524 | 1773.5406 | 1773.16039 | 1773.4247 | 1773.5611 | 1438851.3 | 1773.3228 |
| 1773.4756 | 1324293.3 | 1773.15704 | 1773.536 | 1773.1569 | 1773.3328 | 1773.15707 |
| 1448688.3 | 1773.15135 | 1773.5486 | 1423521.3 | 1448753.3 | 1422042.3 | 1773.5252 |
| 1773.3451 | 1773.3378 | 1773.14868 | 1773.1579 | 1773.338 | 1773.3435 | 1773.4836 |
| 1773.2552 | 1402588.3 | 1354181.3 | 1427193.3 | 1773.15655 | 1455275.3 | 1773.5098 |
| 1324231.3 | 1773.14875 | 1773.15094 | 1773.15925 | 1773.15975 | 1773.15986 | 1448806.3 |
| 1448604.3 | 1773.14933 | 1773.3237 | 1773.15252 | 1773.3474 | 1773.5338 | 1448548.3 |
| 1397868.3 | 1773.4998 | 1773.15333 | 1773.27 | 1773.5706 | 1773.3175 | 1295727.3 |
| 1773.15581 | 1773.5625 | 1773.2515 | 1773.5594 | 1773.5504 | 1773.5442 | 1773.5549 |
| 1773.3048 | 1773.4861 | 1324284.3 | 1773.15813 | 1773.15863 | 1773.15444 | 1773.14784 |
| 1773.5413 | 1448720.3 | 1773.15051 | 1773.2939 | 1773.504 | 1773.15364 | 1773.14776 |
| 1773.522 | 1773.15107 | 1448707.3 | 1773.15551 | 1773.2816 | 1773.356 | 1773.3176 |

**Table 6.** Strains that are used for this experiment

| | | | | | | |
|---|---|---|---|---|---|---|
| 1773.15742 | 1773.15174 | 1773.15599 | 1773.2993 | 1295744.3 | 1417016.3 | 1773.3115 |
| 1400891.3 | 1773.15442 | 1773.15831 | 1397856.3 | 1773.5376 | 1773.5686 | 1773.554 |
| 1448405.3 | 1448555.3 | 1773.15286 | 1448769.3 | 1773.3245 | 1427181.3 | 1773.1542 |
| 1773.15525 | 1773.15351 | 1773.3332 | 1448587.3 | 1773.15557 | 1773.5069 | 1773.3478 |
| 1773.3001 | 1773.15462 | 1773.14845 | 1773.326 | 1773.15082 | 1773.15485 | 1773.15871 |
| 1773.15122 | 1773.15771 | 1773.2895 | 1773.3509 | 1773.5723 | 1773.3398 | 1773.301 |
| 1773.336 | 1773.5455 | 1773.3533 | 1773.3393 | 1418255.3 | 1773.14807 | 1400880.3 |
| 1773.3211 | 1773.15941 | 1773.16086 | 1773.5558 | 1773.3204 | 1773.15006 | 1773.2499 |
| 1773.5572 | 1773.15275 | 1773.4212 | 1773.4206 | 1773.3055 | 1773.2448 | 1773.5207 |
| 1773.3202 | 1773.15121 | 1773.15962 | 1773.4234 | 1773.15463 | 1773.15151 | 1773.294 |
| 1773.15347 | 1773.3529 | 1448527.3 | 1773.2475 | 1773.15988 | 1773.3029 | 1422052.3 |
| 1773.5103 | 1773.2807 | 1773.15272 | 1773.3107 | 1773.3152 | 1773.5144 | 1773.2887 |
| 1773.15024 | 1773.16058 | 1448715.3 | 1773.15063 | 1773.14989 | 1773.15608 | 1773.2564 |
| 1773.3527 | 1773.15246 | 1773.2463 | 1773.5597 | 1773.5282 | 1773.1555 | 1773.15073 |
| 1773.16025 | 1773.3183 | 1773.16008 | 1427180.3 | 1773.3165 | 1773.1525 | 1773.1576 |
| 1448705.3 | 1773.15546 | 1773.15021 | 1448609.3 | 1773.15593 | 1773.3133 | 1773.4999 |
| 1423575.3 | 1448440.3 | 1773.15847 | 1448442.3 | 1773.15883 | 1773.307 | 1324236.3 |
| 1427270.3 | 1773.16031 | 1773.4222 | 1773.283 | 1773.2972 | 1773.15734 | 1773.15765 |
| 1773.14934 | 1448478.3 | 1773.15429 | 1773.2778 | 1773.5143 | 1773.15761 | 1773.2668 |
| 1402598.3 | 1773.15297 | 1773.2473 | 1773.2889 | 1773.3006 | 1397854.3 | 1773.2694 |
| 1773.2568 | 1773.5658 | 1773.16033 | 1773.263 | 1773.3355 | 1773.15241 | 1773.5685 |
| 1400908.3 | 1773.4249 | 1448814.3 | 1448498.3 | 1773.15423 | 1773.2525 | 1438892.3 |
| 1324245.3 | 1773.5616 | 1773.5099 | 1773.15673 | 1447461.3 | 1324285.3 | 1448584.3 |
| 1438852.3 | 1773.14769 | 1773.15388 | 1773.3109 | 1773.14849 | 1773.15179 | 1773.1519 |
| 1773.1596 | 1773.2834 | 1773.3622 | 1773.3437 | 1773.5927 | 1397931.3 | 1773.15244 |
| 1400883.3 | 1400934.4 | 1773.2509 | 1773.2878 | 1773.15932 | 1773.14756 | 1773.318 |
| 1773.553 | 1773.1575 | 1773.5298 | 1773.15096 | 1427275.3 | 1773.3431 | 1773.15648 |
| 1773.5141 | 1773.15173 | 1773.5385 | 1773.15919 | 1773.3376 | 1773.2703 | 1773.15242 |
| 1773.5159 | 1773.258 | 1773.4674 | 1773.5513 | 1448649.3 | 1773.15555 | 1448393.3 |
| 1773.2621 | 1402587.3 | 1773.14938 | 1773.15519 | 1773.15691 | 1773.4253 | 1773.3076 |
| 1773.505 | 1324261.3 | 1773.14869 | 1773.5514 | 1773.2699 | 1448401.3 | 1448725.3 |
| 1773.15795 | 1773.2769 | 1448484.3 | 1773.14805 | 1773.5589 | 1773.1511 | 1455270.3 |
| 1773.5328 | 1773.1518 | 1773.15881 | 1773.2533 | 1427273.3 | 1773.2527 | 1448461.3 |
| 1773.16045 | 1773.3061 | 1773.3097 | 1773.15088 | 1773.5079 | 1773.3125 | 1773.5003 |
| 1773.372 | 1773.3433 | 1773.5133 | 1773.2774 | 1423570.3 | 1448488.3 | 1773.5475 |
| 1447514.3 | 1773.2641 | 1400882.3 | 1773.5116 | 1448654.3 | 33894.6 | 1773.3035 |
| 1773.2957 | 1447464.3 | 1773.14837 | 1773.14932 | 1773.364 | 1408941.4 | 1773.2491 |
| 1773.305 | 1422026.3 | 1773.5104 | 1397927.3 | 1448798.3 | 1773.5025 | 1408947.4 |
| 1773.15394 | 1773.3505 | 1773.4121 | 1773.15533 | 1773.5883 | 1773.2884 | 1423485.3 |
| 1448576.3 | 1773.303 | 1773.5178 | 1773.2661 | 1427316.3 | 1773.3497 | 1773.14945 |
| 1773.15213 | 1773.15328 | 1773.15611 | 1773.15835 | 1773.16095 | 1773.5464 | 1773.5452 |
| 1773.567 | 1773.3409 | 1400916.3 | 1427264.3 | 1773.5246 | 1400918.3 | 1773.3397 |
| 1773.2455 | 1773.14861 | 1773.2628 | 1773.3264 | 1773.5626 | 1773.16097 | 1773.5451 |
| 1773.14751 | 1455289.3 | 1773.371 | 1773.15634 | 1773.15873 | 1773.5652 | 1773.14936 |
| 1423467.3 | 1448789.3 | 1397906.3 | 1773.304 | 1773.2 | 1773.5014 | 1773.15007 |
| 1448607.3 | 1773.15896 | 1773.274 | 1773.15606 | 1773.3472 | 1773.1509 | 1773.1587 |
| 1773.5019 | 1773.14792 | 1773.15724 | 1773.15901 | 1773.421 | 1447471.3 | 1773.307 |
| 1773.15418 | 1448554.3 | 1773.5128 | 1773.37 | 1773.332 | 1773.15198 | 1773.2902 |
| 1408957.4 | 1773.15154 | 1773.2795 | 1773.3419 | 1773.277 | 1295790.3 | 1773.5263 |
| 1408968.4 | 1447456.3 | 1773.14941 | 1773.15903 | 1427228.3 | 1773.14744 | 1354173.3 |
| 1408920.4 | 1447460.3 | 1773.5266 | 1773.15528 | 1448608.3 | 1773.15945 | 1773.2752 |
| 1427229.3 | 1773.3375 | 1773.5352 | 1400927.3 | 1773.5632 | 1773.319 | 1773.526 |

**Table 6.** Strains that are used for this experiment

| | | | | | | |
|---|---|---|---|---|---|---|
| 1773.15145 | 1773.15576 | 1773.15377 | 1448675.3 | 1773.5726 | 1402586.3 | 1773.2567 |
| 1773.282 | 1773.15933 | 1423529.3 | 1773.15119 | 1773.4215 | 1773.15319 | 1773.15384 |
| 1408952.4 | 1773.5286 | 1773.15343 | 1773.3394 | 1773.14844 | 1773.5944 | 1295776.3 |
| 1773.3379 | 1773.3193 | 1773.15999 | 1438877.3 | 1427282.3 | 1773.15066 | 1773.552 |
| 1408930.4 | 1773.558 | 1423566.3 | 1397917.3 | 1448569.3 | 1773.15276 | 1773.15991 |
| 1773.3463 | 1773.3101 | 1773.5332 | 1773.15382 | 1773.1517 | 1773.15421 | 1773.14946 |
| 1408956.4 | 1773.2441 | 1773.2749 | 1400922.3 | 1408921.4 | 1773.1515 | 1773.3511 |
| 1773.5544 | 1773.14731 | 1773.15446 | 1773.1556 | 1773.2498 | 1773.3371 | 1427303.3 |
| 1773.2973 | 1427233.3 | 1773.5891 | 1448726.3 | 1448717.3 | 1448579.3 | 1773.15665 |
| 1773.468 | 1773.15522 | 1773.3126 | 1773.14866 | 1773.3331 | 1773.5532 | 1448829.3 |
| 1408934.4 | 1773.16057 | 1427251.3 | 1773.14907 | 1773.15294 | 1295771.3 | 1773.15857 |
| 1773.2968 | 1773.2901 | 1773.5138 | 1324294.3 | 1773.5498 | 1773.15592 | 1773.15719 |
| 1773.5694 | 1773.2444 | 1438873.3 | 1773.4038 | 1773.5344 | 1773.5297 | 1773.339 |
| 1448490.3 | 1773.2866 | 1448599.3 | 1448733.3 | 1773.2981 | 1773.15311 | 1438878.3 |
| 1773.3301 | 1448492.3 | 1447448.3 | 1448718.3 | 1773.1504 | 1773.5151 | 1773.15631 |
| 1773.302 | 1773.2716 | 1397859.3 | 1773.279 | 1447518.3 | 1773.5106 | 1773.15392 |
| 1773.3217 | 1448563.3 | 1773.15133 | 1773.3033 | 1354158.3 | 1773.2442 | 1773.15305 |
| 1422072.3 | 1397902.3 | 1455286.3 | 1773.5672 | 1773.2496 | 1773.269 | 1773.2758 |
| 1324274.3 | 1773.5362 | 1773.15289 | 1773.15124 | 1448546.3 | 1400888.3 | 1773.14959 |
| 1773.2767 | 1455272.3 | 1773.5185 | 1773.15693 | 1427215.3 | 1448647.3 | 1427243.3 |
| 1773.15753 | 1773.2773 | 1773.5456 | 1773.15979 | 1773.1558 | 1773.15431 | 1773.5121 |
| 1438859.3 | 1448817.3 | 1773.15221 | 1773.4233 | 1773.4864 | 1773.5449 | 1773.3341 |
| 1773.14927 | 1773.16035 | 1773.2711 | 1773.291 | 1324299.3 | 1773.42 | 1773.15927 |
| 1417017.3 | 1773.5339 | 1773.2868 | 1455288.3 | 1773.15131 | 1773.2898 | 1427213.3 |
| 1773.5536 | 1773.4181 | 1773.5325 | 1448464.3 | 78331.98 | 1408962.4 | 1773.5651 |
| 1773.2591 | 1773.3367 | 1773.15526 | 1773.3004 | 1773.14759 | 1773.2915 | 1438879.3 |
| 1773.2859 | 1773.3208 | 1773.3521 | 1773.5052 | 1773.15824 | 1773.15584 | 1773.14948 |
| 1773.5377 | 1773.2574 | 1773.15816 | 1773.14912 | 1773.2663 | 1773.29 | 1773.5373 |
| 1773.271 | 1448635.3 | 1773.15773 | 1773.268 | 1773.15791 | 1773.5709 | 1773.3525 |
| 1295766.3 | 1421921.3 | 1324279.3 | 1773.14821 | 1773.15674 | 1400887.3 | 1773.2636 |
| 1773.3148 | 1773.341 | 1773.5419 | 1773.4262 | 1354182.3 | 1773.5506 | 1773.3468 |
| 1423503.3 | 1773.15752 | 1773.3366 | 1397914.3 | 1773.5375 | 1773.15507 | 1773.2715 |
| 1773.3515 | 1354130.3 | 1427302.3 | 1773.15111 | 1421954.3 | 1773.52 | 1773.15622 |
| 1773.15381 | 1773.15627 | 1773.15358 | 1773.315 | 1773.15146 | 1773.16022 | 1773.3338 |
| 1773.15467 | 1773.15003 | 1773.15105 | 1773.2794 | 1773.5463 | 1773.15089 | 1773.15501 |
| 1773.3466 | 1773.14749 | 1773.2502 | 1773.3059 | 1427308.3 | 1773.15702 | 1773.1526 |
| 1773.3306 | 1773.16068 | 1408906.4 | 1421986.3 | 1773.155 | 1423512.3 | 1448517.3 |
| 1773.5677 | 1421928.3 | 1773.5361 | 1773.14795 | 1408915.4 | 1354129.3 | 1354179.3 |
| 1422008.3 | 1773.5228 | 1773.4852 | 1422087.3 | 1773.2549 | 1447443.3 | 1427199.3 |
| 1448702.3 | 1773.15875 | 1422065.3 | 1448679.3 | 1427226.3 | 1773.548 | 1773.3349 |
| 1773.2583 | 1773.3161 | 1773.5637 | 1773.15921 | 1447480.3 | 1773.15018 | 1773.1478 |
| 1400881.3 | 1773.3377 | 1773.2695 | 1773.3943 | 1773.5525 | 1773.14815 | 1773.15395 |
| 1773.2732 | 1295745.3 | 1773.15352 | 1773.2954 | 1773.342 | 1773.4205 | 1773.5468 |
| 1773.15353 | 1773.1544 | 1773.3429 | 1354164.3 | 1773.3252 | 1773.15571 | 1773.2921 |
| 1773.5916 | 1773.3038 | 1773.15493 | 1448411.3 | 1773.2994 | 1773.16014 | 1773.5154 |
| 1455295.3 | 1773.15676 | 1773.3075 | 1773.2814 | 1773.285 | 1773.2813 | 1773.5693 |
| 1773.3464 | 1773.15996 | 1455264.3 | 1773.14785 | 1773.2445 | 1773.14761 | 1773.2688 |
| 1324298.3 | 1448453.3 | 1773.15189 | 1427305.3 | 1773.15637 | 1324265.3 | 1773.2817 |
| 1773.345 | 1773.3496 | 1773.5537 | 1773.5664 | 1400926.3 | 1773.5369 | 1773.3247 |
| 1773.4867 | 1422013.3 | 1448780.3 | 1773.15181 | 1448729.3 | 1773.1502 | 1400904.3 |
| 1773.2551 | 1773.14733 | 1773.5512 | 1773.15157 | 1448458.3 | 1773.4226 | 1773.1548 |
| 1773.5048 | 1773.5137 | 1447463.3 | 1427311.3 | 1773.5509 | 1295739.3 | 1773.3438 |

**Table 6.** Strains that are used for this experiment

| | | | | | | |
|---|---|---|---|---|---|---|
| 1773.1591 | 1773.1598 | 1773.3145 | 1773.4227 | 1773.5012 | 1773.3234 | 1427205.3 |
| 1354122.3 | 1773.3283 | 1773.1522 | 1773.14847 | 1773.1549 | 1773.2522 | 1773.3299 |
| 1773.15645 | 1773.15741 | 1448625.3 | 1295759.3 | 1773.14895 | 1773.3156 | 1418252.3 |
| 1397928.3 | 1448412.3 | 1448698.3 | 1773.5023 | 1773.14757 | 1773.15509 | 1397884.3 |
| 1423436.3 | 1423558.3 | 1773.14789 | 1773.351 | 1408942.4 | 1773.3207 | 1448636.3 |
| 1400939.3 | 1448568.3 | 1773.2696 | 1773.3461 | 1400872.3 | 1423508.3 | 1324295.3 |
| 1448650.3 | 1773.551 | 1397858.3 | 1773.3226 | 1773.5412 | 1421992.3 | 1455290.3 |
| 1773.3276 | 1408911.4 | 1448606.3 | 1354136.3 | 1354133.3 | 1773.15002 | 1427188.3 |
| 1448459.3 | 1773.5427 | 1773.3318 | 1773.4612 | 1773.4857 | 1773.5233 | 1448559.3 |
| 1773.5728 | 1773.3727 | 1773.3123 | 1773.3488 | 1773.314 | 1295796.3 | 1773.15552 |
| 1773.5167 | 1397867.3 | 1773.3412 | 1773.15192 | 1773.2483 | 1773.2958 | 1773.3064 |
| 1773.5495 | 1773.14877 | 1773.14746 | 1773.14735 | 1773.5247 | 1773.14767 | 1448764.3 |
| 1773.4697 | 1773.15336 | 1773.15661 | 1773.5524 | 1773.14991 | 1773.5683 | 1773.16037 |
| 1773.14888 | 1773.3498 | 1448812.3 | 1773.15214 | 1773.15607 | 1773.3041 | 1773.5215 |
| 1773.4235 | 1773.5587 | 1448745.3 | 1773.16065 | 1773.3178 | 1773.14803 | 1421973.3 |
| 1773.15545 | 1773.5294 | 1773.5326 | 1773.3122 | 1400895.3 | 1773.5642 | 1423517.3 |
| 1773.2558 | 1773.521 | 1773.15758 | 1773.304 | 1773.3637 | 1773.2581 | 1773.3476 |
| 1423440.3 | 1773.15749 | 1448421.3 | 1773.2674 | 1773.3086 | 1773.15898 | 1773.15476 |
| 1773.3073 | 1773.5445 | 1448746.3 | 1773.15372 | 1773.5213 | 1773.3368 | 1773.15516 |
| 1408905.4 | 1773.5013 | 1354110.3 | 1773.2506 | 1773.16076 | 1295767.3 | 1773.4627 |
| 1773.3054 | 1773.2821 | 1773.15186 | 1773.3057 | 1354139.3 | 1773.2824 | 1773.3297 |
| 1773.5542 | 1427320.3 | 1773.278 | 1773.15194 | 1773.2543 | 1773.161 | 1773.2754 |
| 1773.1604 | 1773.15617 | 1773.5617 | 1773.2798 | 1773.2739 | 1773.5522 | 1773.14957 |
| 1773.5232 | 1773.3487 | 1773.3072 | 1408929.4 | 1773.5015 | 1773.5472 | 1773.15247 |
| 1773.15642 | 1773.3469 | 1773.254 | 1773.5574 | 1773.4092 | 1447486.3 | 1448796.3 |
| 1773.3869 | 1773.15128 | 1773.15897 | 1448402.3 | 1773.2672 | 1773.15666 | 1773.3489 |
| 1773.15732 | 1773.2548 | 1408922.4 | 1448468.3 | 1773.5335 | 1324248.3 | 1773.15907 |
| 1773.14904 | 1773.15731 | 1773.3386 | 1773.2684 | 1427183.3 | 1773.5667 | 1773.14741 |
| 1354109.3 | 1773.346 | 1773.15042 | 1773.15915 | 1773.5416 | 1773.2851 | 1773.4678 |
| 1397930.3 | 1773.15796 | 1423531.3 | 1773.2484 | 1773.4601 | 1448433.3 | 1448683.3 |
| 1773.15368 | 1773.2606 | 1773.14812 | 1773.367 | 1447525.3 | 1773.15097 | 1773.15497 |
| 1773.15698 | 1773.2462 | 1773.15385 | 1773.322 | 1773.3576 | 1773.2685 | 1773.3518 |
| 1773.14943 | 1455294.3 | 1773.3233 | 1448398.3 | 1773.5147 | 1773.5092 | 1773.54 |
| 1773.2631 | 1773.5388 | 1448824.3 | 1773.15647 | 1354144.3 | 1448639.3 | 1423446.3 |
| 1773.2653 | 1773.4844 | 1773.2611 | 1773.2885 | 1773.3205 | 1773.15228 | 1773.5287 |
| 1773.15523 | 1773.3124 | 1773.513 | 1773.5699 | 1421983.3 | 1773.2952 | 1324269.3 |
| 1773.14999 | 1773.15963 | 1773.2569 | 1324267.3 | 1773.5183 | 1397919.3 | 1773.5078 |
| 1773.3704 | 1448766.3 | 1773.15825 | 1354176.3 | 1773.274 | 1773.3284 | 1773.15459 |
| 1448681.3 | 1773.16024 | 1773.2723 | 1324233.3 | 1448611.3 | 1448524.3 | 1773.15785 |
| 1773.335 | 1773.4709 | 1421969.3 | 1448467.3 | 1773.2803 | 1448728.3 | 1773.3501 |
| 1773.5584 | 1773.3598 | 1773.292 | 1773.15139 | 1773.5363 | 1773.2797 | 1773.3708 |
| 1423515.3 | 1773.247 | 1773.15046 | 1397939.3 | 1773.2595 | 1773.14914 | 1447506.3 |
| 1773.5551 | 1773.5592 | 1427223.3 | 1773.16009 | 1773.5096 | 1422076.3 | 1773.16005 |
| 1773.2809 | 1773.3345 | 1773.2828 | 1773.2596 | 1427222.3 | 1773.15422 | 1773.2474 |
| 1773.5888 | 1773.4232 | 1773.15895 | 1773.2584 | 1773.5202 | 1773.5679 | 1773.4734 |
| 1397899.3 | 1447482.3 | 1427299.3 | 1773.5669 | 1773.3519 | 1773.15861 | 1773.5497 |
| 1773.2454 | 1773.15885 | 1773.16046 | 1773.5382 | 1773.1593 | 1448823.3 | 1773.5717 |
| 1773.15468 | 1448477.3 | 1773.5428 | 1773.3315 | 1773.3197 | 1448659.3 | 1438836.3 |
| 1773.5168 | 1427301.3 | 1773.15404 | 1773.269 | 1773.5037 | 1773.4263 | 1773.3013 |
| 1773.3235 | 1773.15913 | 1773.15071 | 1773.15101 | 1427214.3 | 1773.15482 | 1773.5517 |
| 1773.15004 | 1773.16103 | 1773.15044 | 1773.15515 | 1773.5907 | 1427280.3 | 1773.3751 |
| 1773.5489 | 1773.357 | 1773.5704 | 1422068.3 | 1773.5153 | 1324230.3 | 1773.3387 |

**Table 6.** Strains that are used for this experiment

| | | | | | | |
|---|---|---|---|---|---|---|
| 1773.2842 | 1773.5477 | 1448721.3 | 1773.3499 | 1773.2943 | 1448436.3 | 1773.3279 |
| 1773.4846 | 1773.3363 | 1773.3321 | 1354124.3 | 1448777.3 | 1773.5368 | 1773.15415 |
| 1773.2464 | 1773.2932 | 1773.15838 | 1448571.3 | 1773.3401 | 1354166.3 | 1295742.3 |
| 1773.15043 | 1423545.3 | 1448773.3 | 1448395.3 | 1354128.3 | 1773.3749 | 1773.5705 |
| 1773.5136 | 1773.5665 | 1448564.3 | 1773.4259 | 1448535.3 | 1427317.3 | 1448759.3 |
| 1773.2516 | 1448665.3 | 1397873.3 | 1267363.3 | 1773.5334 | 1773.1529 | 1773.15303 |
| 1773.5527 | 1773.509 | 1773.15316 | 1448506.3 | 1773.3422 | 1773.15563 | 1773.3118 |
| 1773.3198 | 1773.3222 | 1773.2912 | 1448714.3 | 1448748.3 | 1448664.3 | 1448435.3 |
| 1773.14824 | 1773.15997 | 1448736.3 | 1773.3108 | 1773.3257 | 1773.3423 | 1427276.3 |
| 1773.14774 | 1448828.3 | 1773.545 | 1448751.3 | 1773.16007 | 1773.3 | 1773.3134 |
| 1773.15653 | 1773.2819 | 1773.3309 | 1773.3098 | 1773.15524 | 1773.16084 | 1427237.3 |
| 1773.5091 | 1447437.3 | 1773.15296 | 1773.2607 | 1773.15041 | 1773.5071 | 1773.2804 |
| 1773.5021 | 1773.3298 | 1354141.3 | 1773.4997 | 1438834.3 | 1773.549 | 1773.5582 |
| 1773.3407 | 1423435.3 | 1773.15078 | 1773.15649 | 1773.14753 | 1773.2726 | 1773.14839 |
| 1773.15232 | 1773.15367 | 1773.1523 | 1455307.3 | 1773.5639 | 1773.15514 | 1773.2447 |
| 1773.2625 | 1773.3262 | 1773.5462 | 1427260.3 | 1423562.3 | 1773.3291 | 1773.5336 |
| 1773.2748 | 1447508.3 | 1773.2903 | 1773.3747 | 1773.5599 | 1773.5716 | 1773.15596 |
| 1773.4231 | 1773.15317 | 1773.2755 | 1773.2919 | 1773.2679 | 1417030.3 | 1427288.3 |
| 1773.3159 | 1773.344 | 1773.15565 | 1773.1585 | 1773.555 | 1427285.3 | 1773.15628 |
| 1773.15865 | 1773.15405 | 1773.354 | 1324301.3 | 1447505.3 | 1773.2675 | 1400899.3 |
| 1448836.3 | 1773.2705 | 1773.15716 | 1773.14773 | 1773.14972 | 1773.15669 | 1773.15472 |
| 1773.16085 | 1773.15349 | 1773.2976 | 1773.3192 | 1773.3416 | 1427298.3 | 1423463.3 |
| 1773.3514 | 1773.2937 | 1773.4859 | 1400889.3 | 1324273.3 | 1773.14798 | 1773.15326 |
| 1423448.3 | 1773.5689 | 1773.4848 | 1773.15325 | 1773.2449 | 1773.2914 | 1773.2977 |
| 1397862.3 | 1773.2801 | 1773.1588 | 1773.3304 | 1773.15012 | 1773.543 | 1423501.3 |
| 1773.2856 | 1448666.3 | 1773.15483 | 1448667.3 | 1423489.3 | 1427283.3 | 1773.3987 |
| 1773.5581 | 1455312.3 | 1773.15416 | 1773.323 | 1773.5485 | 1773.15832 | 1773.5605 |
| 1773.5321 | 1773.15473 | 1773.3441 | 1427259.3 | 1773.5209 | 1773.5565 | 1773.5304 |
| 1773.14976 | 1773.15893 | 1773.2883 | 1448514.3 | 1773.5894 | 1448550.3 | 1773.2659 |
| 1773.15029 | 1773.1561 | 1773.5505 | 1773.5305 | 1773.547 | 1448522.3 | 1773.3277 |
| 1400877.3 | 1408945.4 | 1773.5395 | 1448770.3 | 1773.5277 | 1773.15398 | 1421990.3 |
| 1773.3434 | 1448508.3 | 1773.14747 | 1773.2592 | 1427197.3 | 1773.3239 | 1448804.3 |
| 1773.16099 | 1773.3388 | 1421947.3 | 1397940.3 | 1438856.3 | 1773.3327 | 1773.3014 |
| 1773.14942 | 1773.15234 | 1773.2917 | 1773.3009 | 1773.15694 | 1773.5027 | 1773.15474 |
| 1324291.3 | 1447470.3 | 1773.3425 | 1773.2823 | 1773.3456 | 1773.4849 | 1773.1497 |
| 1773.5474 | 1773.565 | 1773.5585 | 1773.15387 | 1773.4238 | 1427241.3 | 1773.5436 |
| 1447495.3 | 1773.339 | 1448605.3 | 1427253.3 | 1773.15215 | 1773.3365 | 1773.14987 |
| 1773.15789 | 1438874.3 | 1773.3167 | 1773.15644 | 1447524.3 | 1448594.3 | 1773.541 |
| 1773.4256 | 1773.3405 | 1423551.3 | 1773.5195 | 1773.3373 | 1397888.3 | 1408913.4 |
| 1773.15064 | 1773.15916 | 1773.3773 | 1773.15797 | 1773.2588 | 1773.3352 | 1773.15153 |
| 1773.15505 | 1773.324 | 1773.15498 | 1773.2951 | 1773.3495 | 1324262.3 | 1417028.3 |
| 1773.5044 | 1773.3141 | 1427318.3 | 1408971.4 | 1773.15783 | 1773.561 | 1773.2644 |
| 1773.4005 | 1427231.3 | 1773.5409 | 1438850.3 | 1773.16001 | 1447492.3 | 1773.2925 |
| 1448390.3 | 1773.14917 | 1773.2702 | 1773.15245 | 1773.15259 | 1773.15323 | 1773.15158 |
| 1423479.3 | 1773.5598 | 1773.15948 | 1773.15574 | 1773.299 | 1773.348 | 1427277.3 |
| 1773.15879 | 1427287.3 | 1773.3027 | 1295805.3 | 1773.15162 | 1773.3214 | 1448797.3 |
| 1773.3063 | 1773.15743 | 1408973.4 | 1773.3482 | 1773.15177 | 1773.5494 | 1773.15746 |
| 1773.539 | 1295800.3 | 1773.3219 | 1773.15248 | 1773.14734 | 1427295.3 | 1423552.3 |
| 1773.5158 | 1773.15947 | 1448802.3 | 1773.4251 | 1773.5275 | 1422059.3 | 1448710.3 |
| 1448723.3 | 1354180.3 | 1773.15894 | 1773.15542 | 1773.15799 | 1773.3005 | 1408949.4 |
| 1773.4217 | 1773.5064 | 1773.5444 | 1773.1602 | 1773.3295 | 1773.16083 | 1773.5603 |
| 1324239.3 | 1773.16093 | 1448653.3 | 1773.3361 | 1773.5264 | 1427242.3 | 1773.3281 |

**Table 6.** Strains that are used for this experiment

| | | | | | | |
|---|---|---|---|---|---|---|
| 1773.318 | 1400885.3 | 1773.5446 | 1773.15426 | 1773.1539 | 1773.2534 | 1773.1483 |
| 1773.15492 | 1773.15502 | 1773.5248 | 1773.14952 | 1773.279 | 1773.336 | 1773.352 |
| 1448831.3 | 1773.14924 | 1773.15561 | 1773.3319 | 1773.4618 | 1773.3224 | 1773.15864 |
| 1773.361 | 1773.1499 | 1773.2692 | 1773.5915 | 1455302.3 | 1295757.3 | 1773.3188 |
| 1448657.3 | 1448737.3 | 1773.2518 | 1448830.3 | 1773.5176 | 1773.2603 | 1773.15172 |
| 1354138.3 | 1773.2508 | 1773.2918 | 1773.592 | 1773.5553 | 1773.15262 | 1773.16012 |
| 1773.2676 | 1773.15613 | 1773.15365 | 1324247.3 | 1773.3016 | 1773.283 | 1422024.3 |
| 1773.3081 | 1773.15759 | 1773.3428 | 1773.328 | 1773.15747 | 1773.15792 | 1295764.3 |
| 1773.3294 | 1773.2997 | 1295806.3 | 1773.5684 | 1447468.3 | 1448523.3 | 1773.5288 |
| 1354177.3 | 1773.297 | 1773.15424 | 1773.3263 | 1773.3577 | 1422104.3 | 1773.14771 |
| 1448474.3 | 1773.14862 | 1773.15484 | 1773.5562 | 1773.5112 | 1773.1514 | 1773.15256 |
| 1773.2438 | 1773.5438 | 1773.15093 | 1773.5519 | 1773.15846 | 1421932.3 | 1773.368 |
| 1773.2965 | 1324287.3 | 1773.15602 | 1773.15839 | 1773.5347 | 1447458.3 | 1773.15451 |
| 1773.252 | 1773.2466 | 1773.15255 | 1773.2472 | 1773.15187 | 1773.2689 | 1773.2931 |
| 1773.2681 | 1773.15972 | 1448685.3 | 1773.15763 | 1408951.4 | 1773.5337 | 1773.14752 |
| 1423518.3 | 1773.5272 | 1773.3213 | 1324221.3 | 1773.15087 | 1773.5314 | 1455268.3 |
| 1402602.3 | 1421945.3 | 1773.4698 | 1427271.3 | 1773.267 | 1448774.3 | 1773.2936 |
| 1397903.3 | 1773.15612 | 1773.16021 | 1773.5357 | 1773.15878 | 1438884.3 | 1773.15099 |
| 1773.14997 | 1408914.4 | 1773.5402 | 1448760.3 | 1773.15703 | 1773.2544 | 1773.533 |
| 1773.16027 | 1773.5646 | 1773.2573 | 1438846.3 | 1354134.3 | 1423548.3 | 1773.15287 |
| 1773.2697 | 1773.5688 | 1773.5552 | 1295774.3 | 1773.5591 | 1773.2922 | 1773.15386 |
| 1773.2896 | 1773.3232 | 1324238.3 | 1423470.3 | 1773.15624 | 1773.3836 | 1773.5643 |
| 1773.4017 | 1295760.3 | 1773.2808 | 1773.295 | 1773.319 | 1773.3323 | 1773.5727 |
| 1773.14928 | 1448644.3 | 1448750.3 | 1773.2576 | 1773.3344 | 1773.5065 | 1773.266 |
| 1773.3194 | 1773.1607 | 1773.2886 | 1448466.3 | 1773.16062 | 1773.5082 | 1773.5633 |
| 1773.15837 | 1773.2519 | 1773.5424 | 1773.5094 | 1773.2953 | 1423495.3 | 1773.15425 |
| 1448460.3 | 1773.5134 | 1773.5236 | 1773.15084 | 1773.5518 | 1773.3017 | 1448449.3 |
| 1773.312 | 1773.5703 | 1438838.3 | 1448485.3 | 1773.15889 | 1448602.3 | 1422085.3 |
| 1773.5674 | 1324259.3 | 1773.15609 | 1773.5547 | 1773.3443 | 1773.3095 | 1427186.3 |
| 1773.15077 | 1447450.3 | 1773.5193 | 1438890.3 | 1773.5439 | 1422009.3 | 1324251.3 |
| 1448614.3 | 1773.3504 | 1773.15697 | 1773.15891 | 1773.1609 | 1773.15652 | 1773.3256 |
| 1438847.3 | 1422014.3 | 1397875.3 | 1773.15683 | 1448816.3 | 1773.15587 | 1773.3445 |
| 1447477.3 | 1773.3026 | 1448447.3 | 1354171.3 | 1773.2557 | 1295791.3 | 1448483.3 |
| 1773.15984 | 1773.2811 | 1773.5166 | 1773.5279 | 1773.3242 | 1773.345 | 1773.5482 |
| 1773.15363 | 1773.16063 | 1427239.3 | 1773.3049 | 1354131.3 | 1773.15183 | 1773.562 |
| 1773.5124 | 1773.15448 | 1448792.3 | 1773.15118 | 1773.14919 | 1773.4863 | 1448410.3 |
| 1773.154 | 1773.3453 | 1773.5299 | 1448573.3 | 1773.5653 | 1400928.3 | 1773.5269 |
| 1773.2837 | 1324218.3 | 1773.14925 | 1773.1495 | 1773.15196 | 1773.2546 | 1773.2626 |
| 1448712.3 | 1773.262 | 1354151.3 | 1773.15283 | 1448785.3 | 1773.15399 | 1773.3507 |
| 1773.15911 | 1773.3746 | 1418251.3 | 1773.2645 | 1773.2984 | 1408904.4 | 1427258.3 |
| 1773.4619 | 1773.1586 | 1773.5534 | 1773.1541 | 1422107.3 | 1773.5324 | 1773.311 |
| 1773.3457 | 1773.14883 | 1773.2563 | 1773.5113 | 1427324.3 | 1773.15852 | 1773.3137 |
| 1773.2761 | 1773.2806 | 1773.5503 | 1773.16015 | 1448533.3 | 1773.15488 | 1773.1564 |
| 1773.5687 | 1421962.3 | 1773.5595 | 1773.15754 | 1324222.3 | 1448803.3 | 1773.5072 |
| 1773.5107 | 1422089.3 | 1773.3266 | 1448392.3 | 1773.15441 | 1773.566 | 1773.15115 |
| 1773.3481 | 1773.3494 | 1773.14858 | 1397885.3 | 1773.5271 | 1773.15678 | 1773.3018 |
| 1773.5654 | 1427269.3 | 1773.2734 | 1773.2871 | 1402585.3 | 1397876.3 | 1448456.3 |
| 1773.15083 | 1773.15781 | 1773.15929 | 1773.4239 | 1448526.3 | 1773.2805 | 1773.15548 |
| 1773.15554 | 1773.3143 | 1773.14841 | 1773.3389 | 1422064.3 | 1354192.3 | 1773.15226 |
| 1773.15779 | 1773.15269 | 1773.1489 | 1773.15313 | 1773.148 | 1773.15944 | 1773.2587 |
| 1417008.3 | 1773.5119 | 1773.2929 | 1427190.3 | 1773.15136 | 1773.15696 | 1773.3002 |
| 1448775.3 | 1773.2852 | 1427309.3 | 1773.4189 | 1773.15711 | 1447526.3 | 1427240.3 |

**Table 6.** Strains that are used for this experiment

| | | | | | | |
|---|---|---|---|---|---|---|
| 1773.3117 | 1422095.3 | 1773.5588 | 1448618.3 | 1773.15403 | 1773.3139 | 1773.14891 |
| 1773.16055 | 1773.15776 | 1773.15337 | 1773.14745 | 1773.15206 | 1773.15417 | 1773.15452 |
| 1773.3528 | 1773.544 | 1773.5254 | 1773.3044 | 1773.5188 | 1773.5235 | 1423442.3 |
| 1773.5404 | 1397887.3 | 1773.3218 | 1773.3209 | 1354140.3 | 1773.14809 | 1773.15672 |
| 1427300.3 | 1773.2488 | 1773.5001 | 1354193.3 | 1773.5629 | 1773.5714 | 1448754.3 |
| 1773.14834 | 1448776.3 | 1354150.3 | 1773.15987 | 1773.3781 | 1773.502 | 1324240.3 |
| 1447467.3 | 1773.2487 | 1773.5146 | 1448811.3 | 1773.3191 | 1773.5057 | 1773.5031 |
| 1417031.3 | 1773.317 | 1773.3417 | 1773.508 | 1773.15408 | 1773.5296 | 1421951.3 |
| 1773.15849 | 1773.2793 | 1773.3154 | 1438845.3 | 1448495.3 | 1773.2945 | 1423546.3 |
| 1773.1505 | 1427194.3 | 1448686.3 | 1354108.3 | 1773.14897 | 1773.15108 | 1773.5329 |
| 1773.3112 | 1438858.3 | 1773.14968 | 1773.15301 | 1773.3065 | 1773.15684 | 1773.4254 |
| 1773.15205 | 1408972.3 | 1773.16059 | 1447500.3 | 1773.15453 | 1773.288 | 1773.4237 |
| 1455269.3 | 1773.4242 | 1773.16041 | 1773.5636 | 1773.2782 | 1447453.3 | 1773.5315 |
| 1773.4128 | 1773.15778 | 1773.5892 | 1773.15586 | 1773.2913 | 1773.3099 | 1773.3342 |
| 1773.365 | 1354146.3 | 1408975.4 | 1773.5417 | 1438853.3 | 1773.15147 | 1295765.3 |
| 1773.2605 | 1773.14944 | 1773.15633 | 1773.15949 | 1773.15243 | 1773.5454 | 1423474.3 |
| 1417021.3 | 1773.2556 | 1427212.3 | 1773.14772 | 1448768.3 | 1773.15268 | 1773.5696 |
| 1773.2632 | 1773.2905 | 1773.15855 | 1773.15169 | 1773.25 | 1773.14966 | 1773.1562 |
| 1773.3185 | 1408958.4 | 1773.14768 | 1773.2597 | 1773.14743 | 1773.1484 | 1773.15588 |
| 1773.4145 | 1773.2966 | 1354183.3 | 1354113.3 | 1400914.3 | 1773.15553 | 1773.5645 |
| 1397937.3 | 1773.15439 | 1773.302 | 1447522.3 | 1773.3096 | 1773.14794 | 1773.517 |
| 1397896.3 | 1773.527 | 1773.507 | 1773.5502 | 1773.35 | 1773.3325 | 1423571.3 |
| 1354178.3 | 1427254.3 | 1427208.3 | 1773.4625 | 1773.5397 | 1448561.3 | 1773.14864 |
| 1773.16017 | 1427265.3 | 1773.14971 | 1438864.3 | 1417015.3 | 1773.2865 | 1773.15309 |
| 1773.15756 | 1773.5303 | 1773.5181 | 1773.5189 | 1773.15236 | 1773.5499 | 1773.2536 |
| 1773.3149 | 1448801.3 | 1773.5476 | 1773.15748 | 1267361.3 | 1773.15095 | 1773.5222 |
| 1455287.3 | 1448475.3 | 1773.3448 | 1773.15688 | 1773.1485 | 1773.15819 | 1773.2476 |
| 1773.2585 | 1447475.3 | 1773.15104 | 1773.4213 | 1423462.3 | 1408950.4 | 1427291.3 |
| 1397932.3 | 1773.15955 | 1773.3774 | 1773.5097 | 1773.2717 | 1448396.3 | 1773.2746 |
| 1773.5943 | 1423511.3 | 1773.15288 | 1773.5223 | 1773.2927 | 1773.15175 | 1773.347 |
| 1408953.4 | 1448813.3 | 1773.15125 | 1773.15103 | 1773.1547 | 1448519.3 | 1773.5511 |
| 1773.15854 | 1773.4203 | 1773.5698 | 1773.511 | 1447517.3 | 1773.16092 | 1773.286 |
| 1448708.3 | 1773.15597 | 1773.3322 | 1397894.3 | 1773.14874 | 1773.14902 | 1773.15185 |
| 1773.352 | 1773.2956 | 1324268.3 | 1773.5418 | 1422055.3 | 1773.15432 | 1773.15635 |
| 1773.5431 | 1773.2841 | 1448835.3 | 1773.316 | 1448838.3 | 1354161.3 | 1423537.3 |
| 1773.1574 | 1773.15818 | 1773.4199 | 1448661.3 | 1448426.3 | 1773.14918 | 1773.5004 |
| 1773.15537 | 1773.3068 | 1773.15292 | 1408931.4 | 1448761.3 | 1773.1606 | 1773.5919 |
| 1773.15285 | 1400902.3 | 1773.15438 | 1773.15559 | 1773.259 | 1773.327 | 1773.15239 |
| 1773.5374 | 1773.5074 | 1773.15225 | 1397892.3 | 1773.3286 | 1354189.3 | 1773.14786 |
| 1773.3588 | 1773.14901 | 1773.15677 | 1773.15028 | 1773.5628 | 1773.15253 | 1773.2906 |
| 1773.485 | 1427195.3 | 1773.3173 | 1773.14896 | 1773.5721 | 1448706.3 | 1773.15327 |
| 1773.5559 | 1773.2832 | 1773.16042 | 1324246.3 | 1773.15543 | 1773.2443 | 1773.5273 |
| 1773.15503 | 1773.5673 | 1773.15202 | 1773.5358 | 1773.2666 | 1773.15738 | 1773.5586 |
| 1423527.3 | 1448739.3 | 1773.15341 | 1773.15354 | 1773.2738 | 1448701.3 | 1773.2545 |
| 1773.15137 | 1773.15324 | 1773.5276 | 1448574.3 | 1773.2962 | 1773.5655 | 1773.321 |
| 1773.14995 | 1417014.3 | 1773.2651 | 1773.5086 | 1438862.3 | 1421971.3 | 1773.2988 |
| 1773.15946 | 1773.15558 | 1773.15076 | 1448742.3 | 1448693.3 | 1773.5922 | 1295723.3 |
| 1448450.3 | 1448534.4 | 1773.15757 | 1773.293 | 1773.15402 | 1438872.3 | 1427182.3 |
| 1773.16023 | 1773.2493 | 1447446.3 | 1773.3085 | 1438848.3 | 1773.2875 | 1773.14728 |
| 1295720.3 | 1773.15116 | 1773.2892 | 1438866.3 | 1773.14981 | 1773.15072 | 1773.16049 |
| 1447519.3 | 1773.15556 | 1417019.3 | 1773.4028 | 1448762.3 | 1448619.3 | 1773.4628 |
| 1773.5197 | 1773.15027 | 1773.3278 | 1773.2577 | 1773.16064 | 1324227.3 | 1773.5117 |

**Table 6.** Strains that are used for this experiment

| | | | | | | |
|---|---|---|---|---|---|---|
| 1773.15967 | 1773.15152 | 1773.3196 | 1773.5058 | 1773.15887 | 1427191.3 | 1448638.3 |
| 1773.201 | 1773.15518 | 1448601.3 | 1773.15535 | 1773.3 | 1427306.3 | 1773.5523 |
| 1773.15908 | 1773.2572 | 1448646.3 | 1773.5343 | 1773.2575 | 1773.298 | 1773.15216 |
| 1773.3492 | 1773.4204 | 1397912.3 | 1773.15499 | 1773.2526 | 1773.4224 | 1773.15737 |
| 1773.3427 | 1773.5899 | 1397907.3 | 1773.1563 | 1773.2478 | 1773.15692 | 1773.2561 |
| 1773.3046 | 1773.15803 | 1421975.3 | 1773.15808 | 1773.15922 | 1448481.3 | 1773.15926 |
| 1773.149 | 1773.5526 | 1324256.3 | 1427211.3 | 1324223.3 | 1773.257 | 1773.3078 |
| 1773.5411 | 1773.3395 | 1773.15132 | 1773.5443 | 1417012.3 | 1773.15258 | 1773.15541 |
| 1773.1503 | 1448719.3 | 1773.3362 | 1773.15914 | 1773.2855 | 1455273.3 | 1773.5641 |
| 1773.2785 | 1773.15378 | 1773.2482 | 1421982.3 | 1324283.3 | 1408925.4 | 1773.2934 |
| 1773.276 | 1773.3317 | 1773.15978 | 1773.4163 | 1773.2667 | 1773.16003 | 1423469.3 |
| 1447472.3 | 1773.15204 | 1773.289 | 1773.4626 | 1324292.3 | 1773.5623 | 1427255.3 |
| 1773.2648 | 1773.2598 | 1397921.3 | 1423477.3 | 1773.2862 | 1773.4257 | 1773.2713 |
| 1773.15621 | 1773.15917 | 1773.15646 | 1773.5075 | 1773.5676 | 1418250.3 | 1773.14854 |
| 1773.5394 | 1448591.3 | 1773.14964 | 1773.1552 | 1448432.3 | 1773.2693 | 1773.3032 |
| 1773.14939 | 1448772.3 | 1448781.3 | 1773.542 | 1773.5006 | 1773.15411 | 1773.3261 |
| 1773.287 | 1773.15009 | 1773.5708 | 1448763.3 | 1773.265 | 1773.2963 | 1773.4172 |
| 1773.2961 | 1773.16082 | 1422018.3 | 1773.366 | 1422045.3 | 1773.2602 | 1448834.3 |
| 1773.294 | 1427217.3 | 1423426.3 | 1773.1496 | 1773.15176 | 1773.5203 | 1773.5391 |
| 1423557.3 | 1773.15982 | 1773.16098 | 1773.2532 | 1773.15302 | 1423543.3 | 1773.16051 |
| 1773.159 | 1295758.3 | 1773.2978 | 1773.14969 | 1773.4086 | 1773.3374 | 1773.5129 |
| 1773.512 | 1448454.3 | 1324266.3 | 1773.15166 | 1773.15266 | 1427238.3 | 1455305.3 |
| 1773.153 | 1773.15355 | 1773.2537 | 1773.14825 | 1773.2665 | 1773.1501 | 1773.15845 |
| 1773.202 | 1773.3128 | 1773.33 | 1773.15625 | 1773.3236 | 1773.5359 | 1773.5423 |
| 1773.15681 | 1773.4853 | 1773.5 | 1773.14848 | 1773.3189 | 1773.15075 | 1773.15211 |
| 1773.14851 | 1773.5262 | 1773.5306 | 1418254.3 | 1773.3403 | 1773.14982 | 1773.331 |
| 1773.2529 | 1773.277 | 1422044.3 | 1773.14833 | 1773.2933 | 1448692.3 | 1773.14965 |
| 1773.2617 | 1448837.3 | 1773.14967 | 1773.5243 | 1773.14778 | 1773.5318 | 1773.2594 |
| 1773.5528 | 1773.5533 | 1427248.3 | 1773.5367 | 1438861.3 | 1773.5435 | 1773.15049 |
| 1773.3039 | 1324289.3 | 1324258.3 | 1773.295 | 1447523.3 | 1773.16011 | 1773.5634 |
| 1773.14905 | 1773.3067 | 1773.15068 | 1773.15433 | 1773.1592 | 1448670.3 | 1773.3479 |
| 1773.2654 | 1447507.3 | 1773.5225 | 1447440.3 | 1773.2646 | 1773.2615 | 1773.3669 |
| 1773.5372 | 1773.2538 | 1773.3532 | 1423574.3 | 1773.5619 | 1773.15953 | 1773.15968 |
| 1773.14955 | 1423509.3 | 1773.3146 | 1773.5265 | 1773.1474 | 1438887.3 | 1427177.3 |
| 1422082.3 | 1447484.3 | 1354157.3 | 1773.4407 | 1773.5609 | 1400875.3 | 1448629.3 |
| 1423505.3 | 1773.264 | 1773.3093 | 1773.15815 | 1773.15035 | 1773.366 | 1773.5386 |
| 1773.3273 | 1427244.3 | 1773.3275 | 1773.16078 | | | |
